# A Genetic History of Continuity and Mobility in the Iron Age Central Mediterranean

**DOI:** 10.1101/2022.03.13.483276

**Authors:** Hannah M. Moots, Margaret Antonio, Susanna Sawyer, Jeffrey P. Spence, Victoria Oberreiter, Clemens L. Weiß, Michaela Lucci, Yahia Mehdi Seddik Cherifi, Francesco La Pastina, Francesco Genchi, Elisa Praxmeier, Brina Zagorc, Olivia Cheronot, Kadir T. Özdoğan, Lea Demetz, Selma Amrani, Francesca Candilio, Daniela De Angelis, Gabriella Gasperetti, Daniel Fernandes, Ziyue Gao, Mounir Fantar, Alfredo Coppa, Jonathan K. Pritchard, Ron Pinhasi

## Abstract

The Iron Age was a dynamic period in central Mediterranean history, with the expansion of Greek and Phoenician colonies and the growth of Carthage into the dominant maritime power of the Mediterranean. These events were facilitated by the ease of long-distance travel following major advances in seafaring. We know from the archaeological record that trade goods and materials were moving across great distances in unprecedented quantities, but it is unclear how these patterns correlate with human mobility. To investigate population mobility and interactions directly, we sequenced the genomes of 30 ancient individuals from coastal cities around the central Mediterranean, in Tunisia, Sardinia, and central Italy. We observe a meaningful contribution of autochthonous populations, as well as highly heterogeneous ancestry including many individuals with non-local ancestries from other parts of the Mediterranean region. These results highlight both the role of local populations and the extreme interconnectedness of populations in the Iron Age Mediterranean. By studying these trans-Mediterranean neighbors together, we explore the complex interplay between local continuity and mobility that shaped the Iron Age societies of the central Mediterranean.

## Introduction

The 1st millennium BCE was characterized by a marked increase in mobility in the Mediterranean. Advances in sailing and seafaring allowed for easier and more frequent travel across the open sea, facilitating new networks of interaction - for trade, colonization, and conflict. In this paper, we use data from three key port and coastal cities in the central Mediterranean -- Kerkouane (Tunisia), Sant’Imbenia (Sardinia), and Tarquinia (central Italy) -- to study mobility within this region.

In the late Bronze and early Iron Age (1250 – 800 BCE), Phoenician and Greek city-states established trading ports and colonies across the Mediterranean. Carthage was located at the crossroads of these trans-Mediterranean trade routes and, for 500 years (Fig. 1A), was the center of a trading network spanning much of the central and western Mediterranean. Carthage was the dominant maritime power of the region until Roman Imperial expansion in the final centuries BCE ^1,2^. Across the sea, Etruscan-speaking city-states flourished in central and northern Italy. Their material culture suggests continuity with the preceding Villanovan culture, but there are also strong indications of contact with other cultures, both by sea and land ^3^. Insights into the relationship between these trans-Mediterranean neighbors are limited. Few Punic or Etruscan written sources have survived to the present day, and most of these are short inscriptions ^3,4^. As a result, much of the historical record is filtered through the lens of Greek, Roman and Egyptian sources ^5^. Archaeogenetic research adds a new line of evidence to our understanding of the people and their interactions during this dynamic period.

**Fig. 1.**
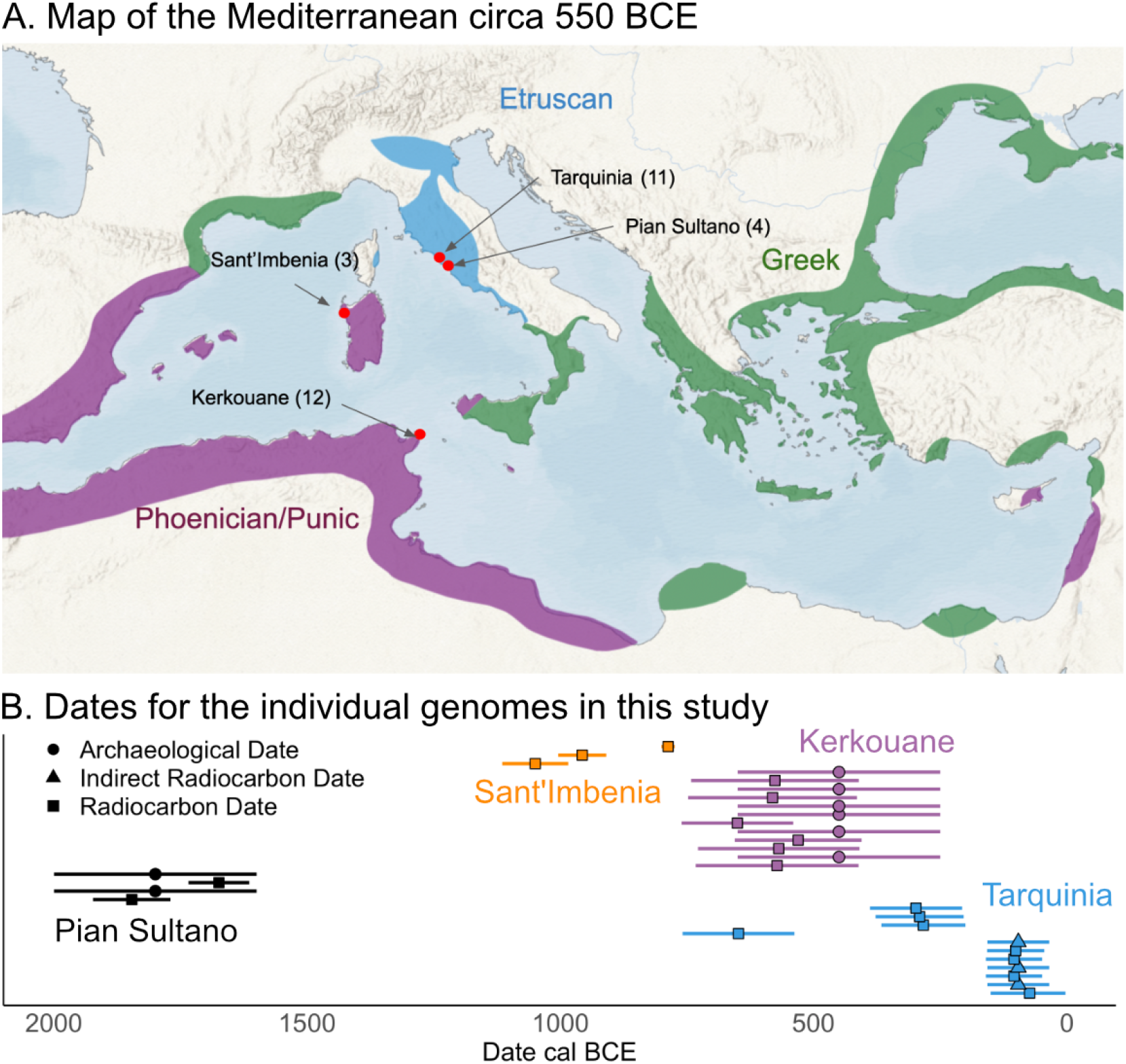
Data overview and relevant geography and chronology. **A**. Locations of the 4 archaeological sites examined here, as well as a map of the areas settled by Phoenician, Greek, and Etruscan speakers by 550 BCE. **B**. Timeline showing the dates for the newly reported individual genomes, with the methods of dating indicated. Indirect radiocarbon dating refers to using the radiocarbon date of a different individual in the same tomb.

Until recently, the genetic makeup of Iron Age Mediterranean populations has been relatively understudied in comparison to earlier periods. New research has begun to shed light on the Iron Age peoples of central Italy and Sardinia ^6–10^ but there is very limited information about the Iron Age North African communities. To date, there has been no whole-genome ancient DNA research on Northwest Africa from this time period, despite the major historical importance of Carthage and the complex history of the region. Mitochondrial DNA from an Iron Age individual from Carthage is the only existing ancient DNA data from the region ^11^.

We illuminate the complicated interplay between continuity and mobility and document extensive genetic exchange across the central Mediterranean which allows us to directly examine relationships between local and diasporic people of the Iron Age. More specifically, we observe heterogeneity at the port cities of the Iron Age central Mediterranean, in accord with historically and archaeologically attested connections. One surprising deviation from this is the absence of individuals with Levantine ancestries at Kerkouane, Sant’Imbenia, and other published Iron Age sites in Sardinia. We also explore whether these mobility patterns varied by sex and examine indications of admixture between individuals of a diverse range of genetic ancestries.

## Results

### Data generation

We generated double-stranded DNA libraries after partial uracil-DNA glycosylase (UDG) treatment. After authentication and screening, libraries were sequenced on an Illumina NovaSeq 6000 sequencing platform to generate whole-genome shotgun data, with an average genome-wide coverage of 1.1x (range: 0.61 - 1.9x). We successfully obtained data from 30 individuals from 4 archaeological sites, shown in Fig. 1A: Kerkouane in Tunisia (n=12), Sant’Imbenia in Sardinia (n=3), and Pian Sultano (n=4) and Tarquinia in central Italy (n=11). In our analyses, the new data are supplemented with 9 previously published Iron Age Sardinians dating from 818- 208 BCE ^6,7^ and 33 additional Iron Age central Italians dating from 963 - 200 BCE ^8,9,12^.

We conducted radiocarbon dating on 19 individuals (Fig. 1B, Dataset S3). These confirm the archaeologically attested dates of use for the Kerkouane necropolis to the mid-Iron Age (650 - 250 BCE), a key period for the growth of Carthage and its role as a primary node in the trade networks of the Mediterranean. In Italy, the individuals from Tarquinia spanned the Iron Age, from the city’s growth in the early Iron Age through its incorporation into the growing Roman Republic in the 3rd century BCE. The burials at Pian Sultano date to the middle Bronze Age, and thus provide important context for central Italian populations before the Iron Age. Individuals at Sant’Imbenia, Sardinia date to the Bronze/Iron Age transition on the island (1115 - 775 cal BCE).

To contextualize these individuals within the genetic landscape of contemporaneous populations, we curated a set of published ancient genomes spanning the Iron Age Mediterranean ^6–8,13–18^. We organized these data into 5 regional groupings: the Italian Peninsula, Sardinia, Northwest Africa/Maghreb, the Levant, and Iberia. To compare with the preceding period, we also curated the available Bronze Age samples for each of these regions ^12,19,20^. For the Maghreb, we included Late Neolithic individuals as there are no published Bronze Age data ^21^. This resulted in a set of 330 reference genomes to contextualize the new data (more information about these and our curation of the metadata can be found in Dataset S2).

### Increased genetic heterogeneity in the Iron Age

We generated a Principal Component Analysis (PCA) reference space using modern populations from around the Mediterranean, Europe, the Middle East, and North Africa ^22^, and projected the Bronze and Iron Age Mediterranean individuals onto this modern reference space (Fig. 2, Fig. 3A). This regional time series approach allows us to make some general observations about changes occurring across these periods. In particular, we observe a marked increase in heterogeneity in the Iron Age. While Bronze Age populations in this region, especially Iberia, had some outlier individuals, they were relatively few in number and sporadic. In the Iron Age, however, there is much more overlap in PCA space between individuals from different regions.

**Fig. 2.**
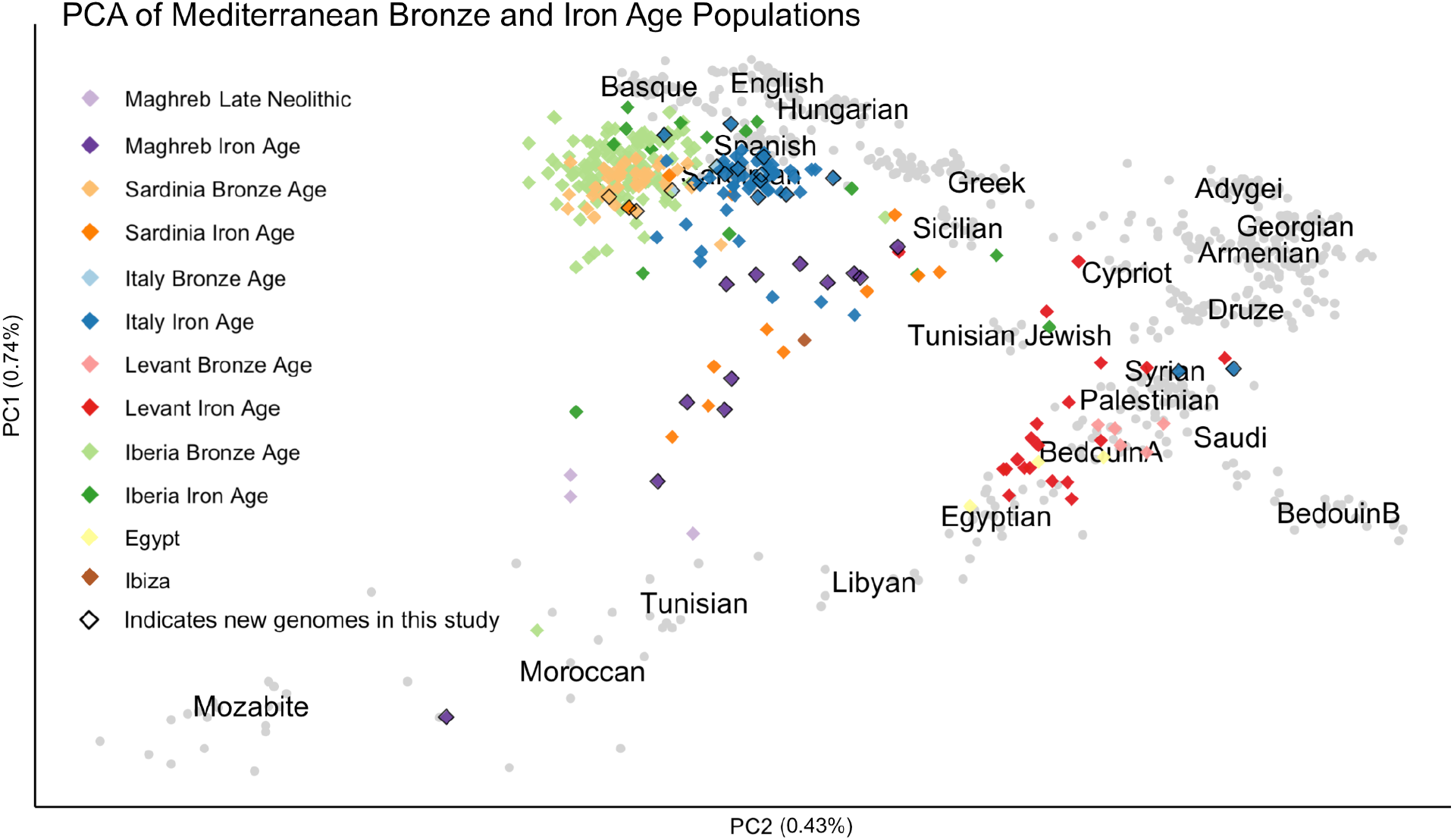
Principal Component Analysis of relevant samples. Newly reported individual genomes (diamond-shaped points) are projected onto a PCA space of present-day individuals (gray points) using smartPCA. Ancient genomes are shown in paired sets of colors by region, with the earlier time period (Bronze Age or Neolithic) represented by the lighter colors and the later time period (Iron Age) represented by the darker colors.

**Fig. 3.**
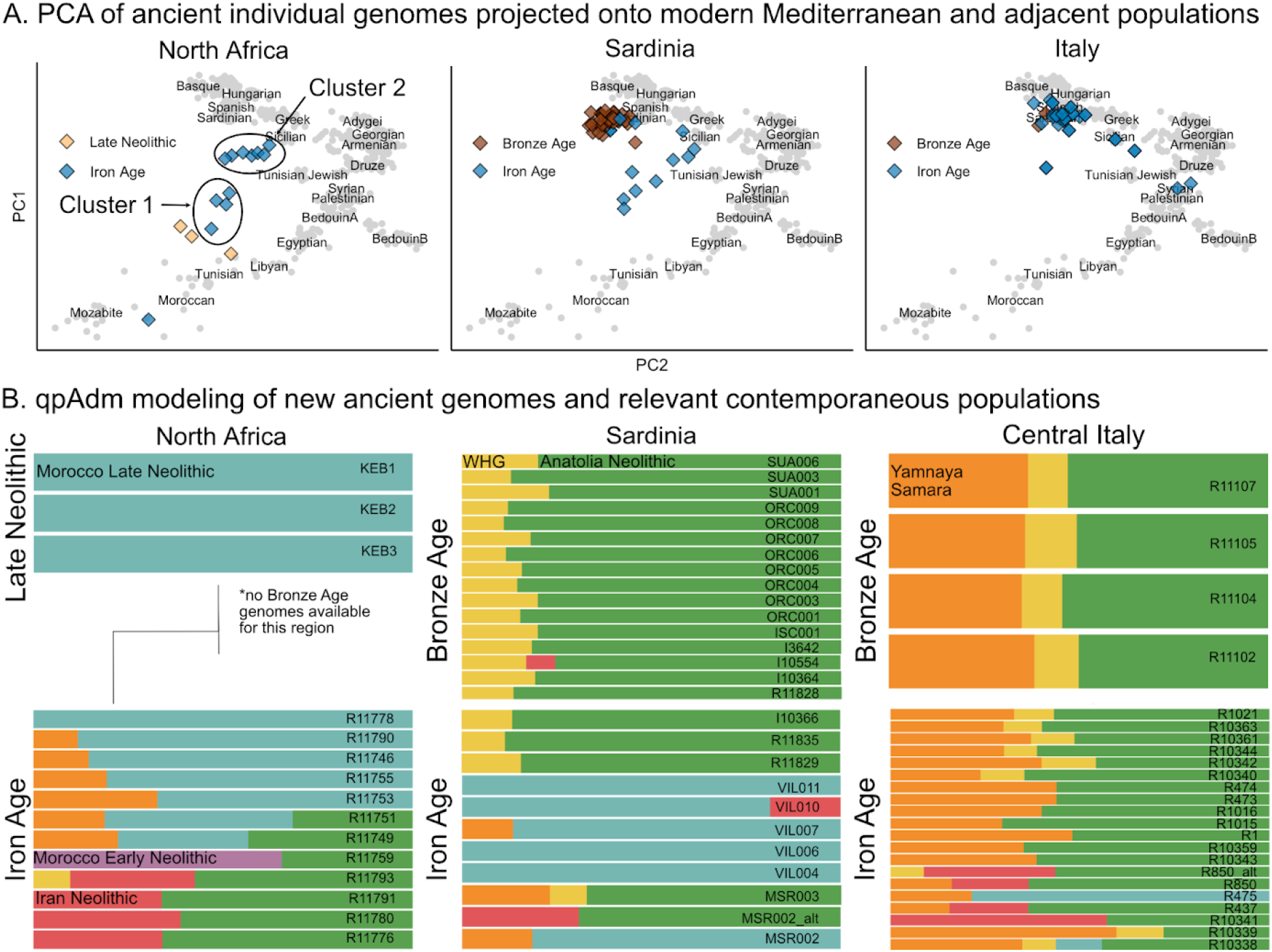
Genetic overview of central Mediterranean populations during the Bronze and Iron Ages. Both PCA and qpAdm illustrate the shift from relative genetic homogeneity in the Bronze Age to heterogeneity in the Iron Age. **A:** PCA of ancient Neolithic, Bronze and Iron Age populations from the Central Mediterranean, projected onto a modern reference space. **B:** qpAdm analyses using a set of distal source populations (Western Hunter-Gatherer (WHG), Yamnaya Samara, Anatolia Neolithic, Iranian Neolithic, Morocco Late Neolithic, and Morocco Early Neolithic).

To further explore these patterns, we modeled the ancestry of the new and published individual genomes from the central Mediterranean with qpAdm admixture modeling, shown in Fig. 3 ^23,24^. We chose a set of distal source populations previously shown to be informative for understanding the diversity of the Mediterranean during this period: Western Hunter-Gatherer (WHG), Yamnaya Samara, Anatolian Neolithic, Iranian Neolithic, and Neolithic farmers from Morocco ^6,8,15^. We next used this same set of source populations to perform pairwise qpWave on all individuals to test whether pairs of individuals form a clade with each other, relative to a set of reference populations (Supplementary Methods). We used 1 - log(p-value) to calculate distances between each pair of individuals and performed clustering (Fig. 5). Given the genetic heterogeneity that characterizes the Iron Age Mediterranean, these groupings identified in qpWave identify genetically similar individuals across regions.

**Fig. 5.**
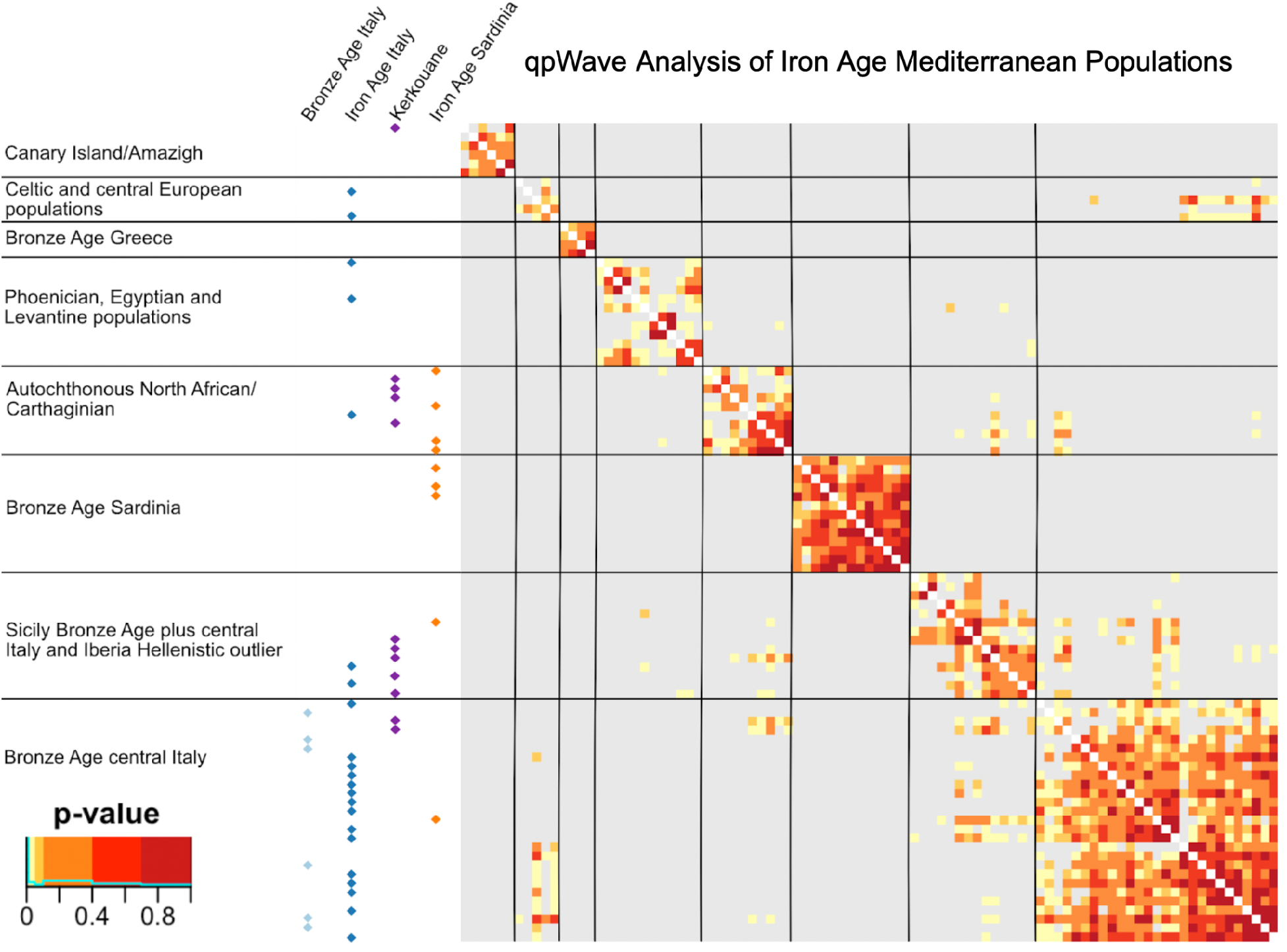
qpWave Analysis of Mediterranean Populations. The plot shows the clustering of Bronze Age and Iron Age genomes from the central Mediterranean with other relevant populations. Heatmap values indicate whether each pair of individuals can be modeled with the same ancestry components in qpAdm in comparison to a set of reference populations. Individual labels are shown in Supplementary Fig. 7.

**Fig. 6.**
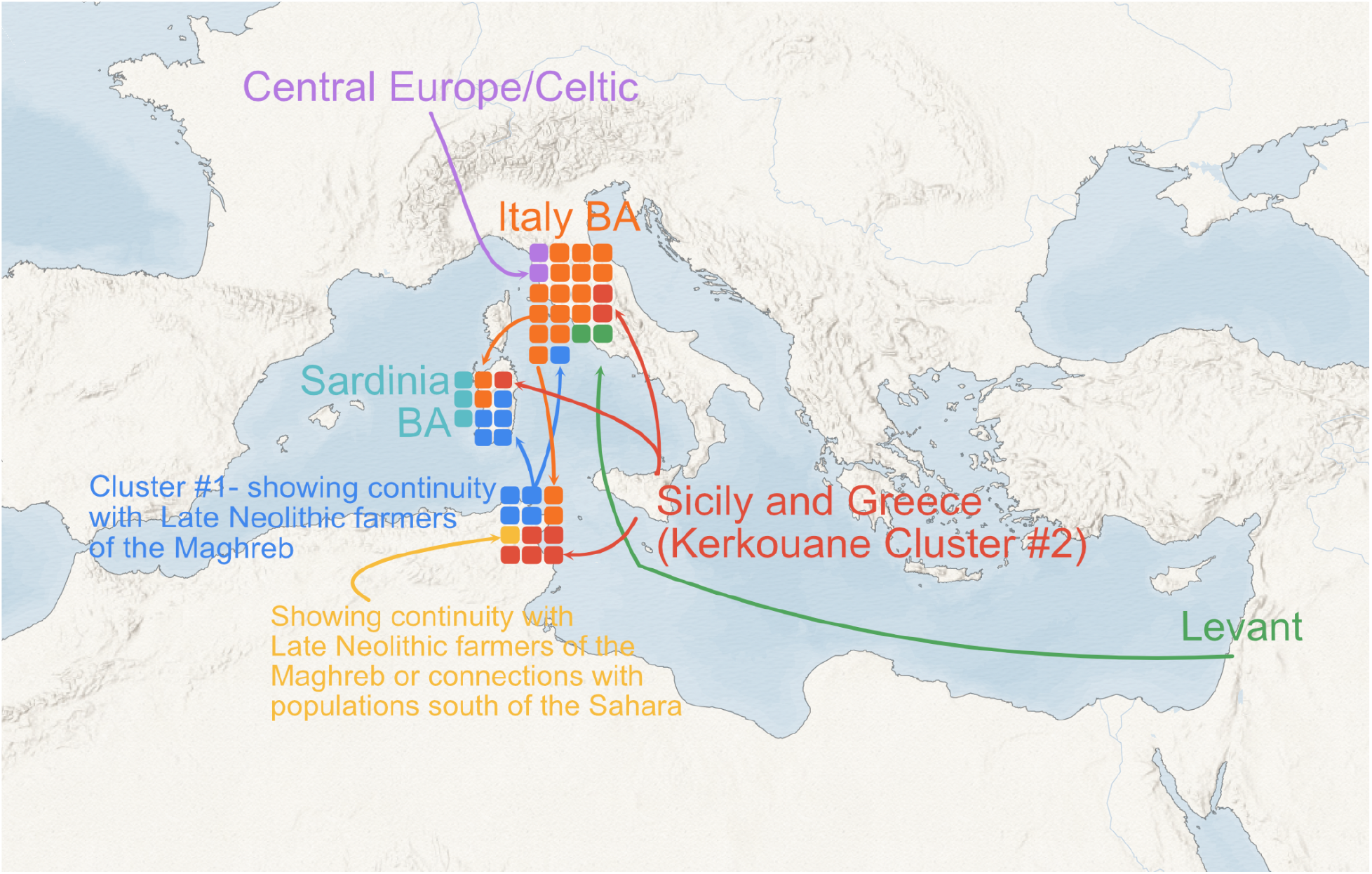
Overview of results. Map showing the memberships of central Mediterranean Iron Age individuals in different qpWave clusters, organized by site. Regional labels and arrows show likely sources of ancestries (the arrows are not intended to indicate specific routes). Colors indicate ancestry clusters as identified by qpWave.

### Characterizing the Genetic Heterogeneity at Kerkouane

Kerkouane is an exceptionally well-preserved town located on Tunisia’s Cap Bon Peninsula and provides one of the best-surviving windows into Carthaginian daily life (M. H. Fantar 2000; H. Fantar 1988; M. H. Fantar 1987; Miles 2010). Originally inhabited from 650 - 250 BCE, the population of Kerkouane is thought to have been around 1,200 with an economy primarily based on the production and export of marine resources from the region, including the production and exportation of garum, salt, and Tyrian purple dye derived from locally harvested Murex sp. shells (M. Fantar 2003). Kerkouane was abandoned after the 1st Punic War between Carthage and Rome and was never re-inhabited or rebuilt by the Romans resulting in the excellent preservation of the original architecture of the town. Scholars have also observed strong Libyan and Greek influences in the material culture recovered at the site (Fantar 1987).

Examining 12 individual genomes from 4 shared tombs, we observe a highly heterogeneous population, projected across the PCA space in Fig. 2 consisting of three primary genetic clusters. One of the genetic groups we identified (labeled “Cluster 1”) includes four individuals who project near preceding (late Neolithic) Maghrebi farmers. We used qpAdm modeling to test whether there was genetic continuity with preceding populations (Fig. 3). One individual, R11778, can be modeled in qpAdm with 100% Morocco Late Neolithic farmer ancestry, while three individuals, R11746, R11755, and R11790, can be modeled predominantly with this component, along with the addition of 15 - 20% Steppe-related ancestry. This suggests that these individuals represent an autochthonous North African population, with some evidence for admixture with individuals of Steppe-related ancestry. Of the 4 individuals in Cluster 1, two have mitochondrial haplogroups which were also identified among the preceding Maghrebi populations (Fregel et al. 20218). R11778 carries haplogroup L3e2bA, which is most common in western Africa, indicating potential trans-Saharan connections during this period ^25^. The biologically male individual in Cluster 1 carries Y-haplogroup R1b, which is associated with Bell Beaker cultures in Europe and common in central Italy at this time and became prevalent in southern Iberia by the Bronze Age ^18^. This supports the possibility that the steppe component observed at Kerkouane may have been introduced through mobility occurring in the central and western Mediterranean, as also suggested by Fernandes et al. 2020. This is also supported by proximal qpAdm modeling, which finds Bell Beaker populations from Iberia and France as suitable proximal sources for this component (Supplementary Table 4).

A second group, labeled “Cluster 2” in PCA (Fig. 3 and Supplementary Fig. 5) and identified in qpWave (Fig. 5), contains seven individuals who cannot be modeled using preceding north African population and are better modeled as genetically similar to Bronze Age Sicilian and central Italian populations, as well as some contemporaneous individuals from the Iberian Greek-speaking colony of Empúries ^15,26^. We cannot know if these individuals with non-local ancestries moved in their lifetime or were part of multi-generational settlement in the region. Of the 7 individuals in the non-local ancestry cluster (Cluster 2), all individuals carry mitochondrial haplogroup HV, H, and I5, all thought to have originated in the Mediterranean region ^27^. All biological males in this cluster carry J2b, which has been reported at high rates in Sardinian and Levantine Bronze Age populations and has been suggested to be associated with the Phoenician expansion. Interestingly, R11790 (in Cluster 1, characterized by genetic continuity with previous populations of the region) also carries HV13 which suggests admixture between the local and diasporic populations at the site.

R11753 and R11791, both from Cluster 2, show strong evidence of inbreeding, with runs of homozygosity (ROH) segments over 50 Mb in length. In both cases, with over 12% of the genome being homozygous (segments > 5Mb in length), the parents were likely 2nd-degree relatives. Consistent with the homozygosity analysis, both individuals also have very low conditional heterozygosity (Fig. S9). While it has been shown that endogamous marriage practices were common in the contemporaneous ancient Greek world ^28^, less is known about such practices in the Carthaginian world, due to fewer surviving written records ^29^.

For R11759, who projects near modern Mozabite and Moroccan populations in PCA space, there were no working distal qpAdm models with the original set of 5 distal source populations (Fig. 5). We replaced Morocco Late Neolithic with Morocco Early Neolithic farmers and a hunter-gatherer individual from Ethiopia from ∼4500 BP ^30^, both of which produced working models. Using competition modeling (where possible sources are rotated to the outgroup), the best model uses ∼70% Morocco Early Neolithic ancestry and ∼30% Anatolia Neolithic (Fig. 3, Supplementary Fig. 5). When compared to other ancient individuals using qpWave analysis (Fig. 5), this individual forms a clade with ancient Canary Island inhabitants thought to be representative of the original founding population ^31^. The Canary Islands were originally settled in the 1st millennium BCE by a population genetically ancestral to today’s populations of North Africa ^32^. This individual carries U6, a common northern African mitochondrial haplogroup.

Based on the study of mitochondrial haplogroups, we observe high rates of mobility, accompanied by admixture (Supplementary Figures S11 and S12). This is consistent with Matisoo-Smith 2018’s finding of heterogeneity in Phoenician-speaking communities at Monte Sirai (Sardinia) and Bey (Lebanon), although the profile of haplogroups observed at Kerkouane differs from those observed at Monte Sirai and Bey ^33^.

### Iron Age Communities in Sardinia and Ibiza

The Iron Age was a dynamic period in Sardinia - initially independent, and later incorporated into the Phoenicio-Punic/Carthaginian trading world and then the Roman empire. Sardinia was a relatively genetically homogenous population through the Bronze Age, with increasing heterogeneity in the Iron Age ^6,7^ and this new genetic information about contemporaneous North Africa allows us to directly explore the connections between Sardinia and North Africa in this period.

The three newly-reported Late Bronze Age/Early Iron Age genomes from Sant’Imbenia (R11828.SG, R11829.SG, R11835.SG; 1115 - 775 cal BCE) show continuity with the preceding Bronze Age Nuragic population. Later individuals (818 - 392 cal BCE) from the site of Villamar (VIL004, VIL006, VIL007, VIL010, VIL011) can be modeled with primarily Morocco Late Neolithic ancestry (Marcus, et al. 2022), similar to the North African cluster from Kerkouane (Cluster 1), as well as a individual from Ibiza, MS10614 ^16^, which mirrors the strong political and trading connections at the time with Carthaginian North Africa. This timing is consistent with a previous estimate from modern genomes that African gene flow into Sardinia occurred 96 generations (∼2750 years ago) ^34^. Contemporaneous individuals (800 - 300 BCE) from the site of Monte Sirai (MSR002, MSR003), cluster with contemporaneous central Italian and Sicilian populations (Fig. 5). Following the Roman annexation of this territory from Carthage in the First Punic War (241 BCE), Roman period individuals from Sardinia project in PCA near contemporaneous individuals from mainland Italy ^6,7^. We observe a shifting genetic profile reflecting the shifting political affiliations of the island through the Iron Age (Supplementary Fig. 3).

### Genetic Evidence for connection across the Central Mediterranean

In Iron Age Italy, consistent with the findings of Posth et al. 2021, we observe a combination of genetic continuity from the Bronze Age population, along with a drastic increase in genetic heterogeneity compared to the Bronze Age. Of the 22 Iron Age central Italian genomes, 12 individuals form a clade in the qpWave and qpAdm analyses (Figs. 4, 5) with Bronze Age central Italian individuals, and thus can be modeled as deriving 100% of their ancestry from Bronze Age central Italians. However, nearly half of the individuals are best modeled in qpWave with ancestry from other parts of the Iron Age Mediterranean world (Fig. 3). Among these, one individual, R475, projects near the clade of autochthonous North African individuals at Kerkouane and can be modeled with the sample distal source populations in qpAdm (∼85% Late Neolithic Moroccan farmer and∼20% steppe-related ancestry). Additionally, four recently published individuals from Tarquinia have northern African ancestry ^9^. Two individuals, R10337 and R10341, appear genetically similar to contemporaneous individuals from the Levant (Figs. 3, 4). In ADMIXTURE modeling, these two individuals lack Steppe-like ancestry, contrasting with other contemporaneous individuals from central Italy. They both have high amounts of Iranian Neolithic ancestry (Supplementary Fig. 4), and may be early instances of the shift towards eastern Mediterranean ancestry of the following Roman imperial period ^8^. This component appears in high levels in the two outlier individuals mentioned above and then in smaller amounts later in the Imperial period population of Rome (Fig. 2, Supplementary Fig. 3). This suggests this ancestry may have instead been introduced through individuals traveling long distances within their own lifetimes, as we know was happening with the establishment of Phoenician and Greek colonies throughout the Mediterranean at this time. In contrast, the appearance of steppe-related ancestry during the Bronze Age in central Italy originally occurred in small amounts, ubiquitously in the population (Saupe et al. 2021), suggesting that this ancestry may have spread gradually through small, local interactions over many generations.

### Iron Age Mobility Shaped the Present-Day Distribution of Ancestry Components of the Region

Across all three locations, we observe continuity in the Iron Age with the preceding population of the region accompanied by the presence of individuals with ancestry from other parts of the Mediterranean world and beyond. While the high heterogeneity at these coastal cities is striking, we were curious if the mobility indicated thereby was limited to port cities or extended into the hinterlands as well. While there is not available ancient data from the rural contexts surrounding these ports, modern data offer insights into whether the ancestries introduced in the 1st millennium BCE were transient (and thus isolated to the port cities) or permanently impacted the population structure of these regions, indicating that this mobility extended beyond the ports, albeit perhaps at lower rates. In qpAdm and ADMIXTURE modeling (Fig. 3, Supplementary Fig. 4), three ancestral components were generally sufficient for modeling the pre-Iron Age populations in North Africa, Sardinia, and Italy, but in the Iron Age, a minimum of five components are needed to model the ancestry profiles of individuals at the sites studied. The Morocco Late Neolithic component, which was predominantly found in North Africa prior to the Iron Age, now appears in central Italy, as well as in individuals from Carthaginian sites across the central and western Mediterranean, such as Ibiza. This component may be part of the genetic signature of Carthage as a trading hub and maritime power in the region. This mobility seems to have contributed to the present-day genetic structure of these populations. Parallel to contemporaneous populations in the Eastern Mediterranean and Southwest Asia ^35^, increasing interconnectivity was also paired with increasing long-distance gene flow patterned by the historical, geopolitical, and even environmental conditions of the time, maintaining population structure in the region ^36^.

## Discussion

By integrating our research questions across the populations of the central Mediterranean, we describe here how increasing rates of mobility across the Mediterranean contributed to the genetic history of the region. The high number of individuals with North African ancestry from central Italy may reflect the close connections between Carthage and the Etruscan-speaking city-states, both through trade and also, at times, as allies facing common adversaries, especially Greek and Roman Imperial expansion, such as at the Battle of Alalia around 535 BCE to oppose Greek expansion in central Italy. In particular, our findings support and add temporal resolution to previous suggestions based on modern and historical data ^34,36,37^ that the Iron Age was a key time for trans-Mediterranean mobility and connectivity between the regions we today call northern Africa and southern Europe. Supporting this, the presence of a number of individuals similar to contemporaneous Italian and Greek populations at Kerkouane suggests a bidirectional movement of people, especially within the central Mediterranean. At both Kerkouane and Tarquinia, we observe that individuals buried together have diverse and geographically distant ancestries (Table 1). Non-local ancestry doesn’t seem to have necessarily resulted in differential treatment in funerary celebrations.

**Table 1.**
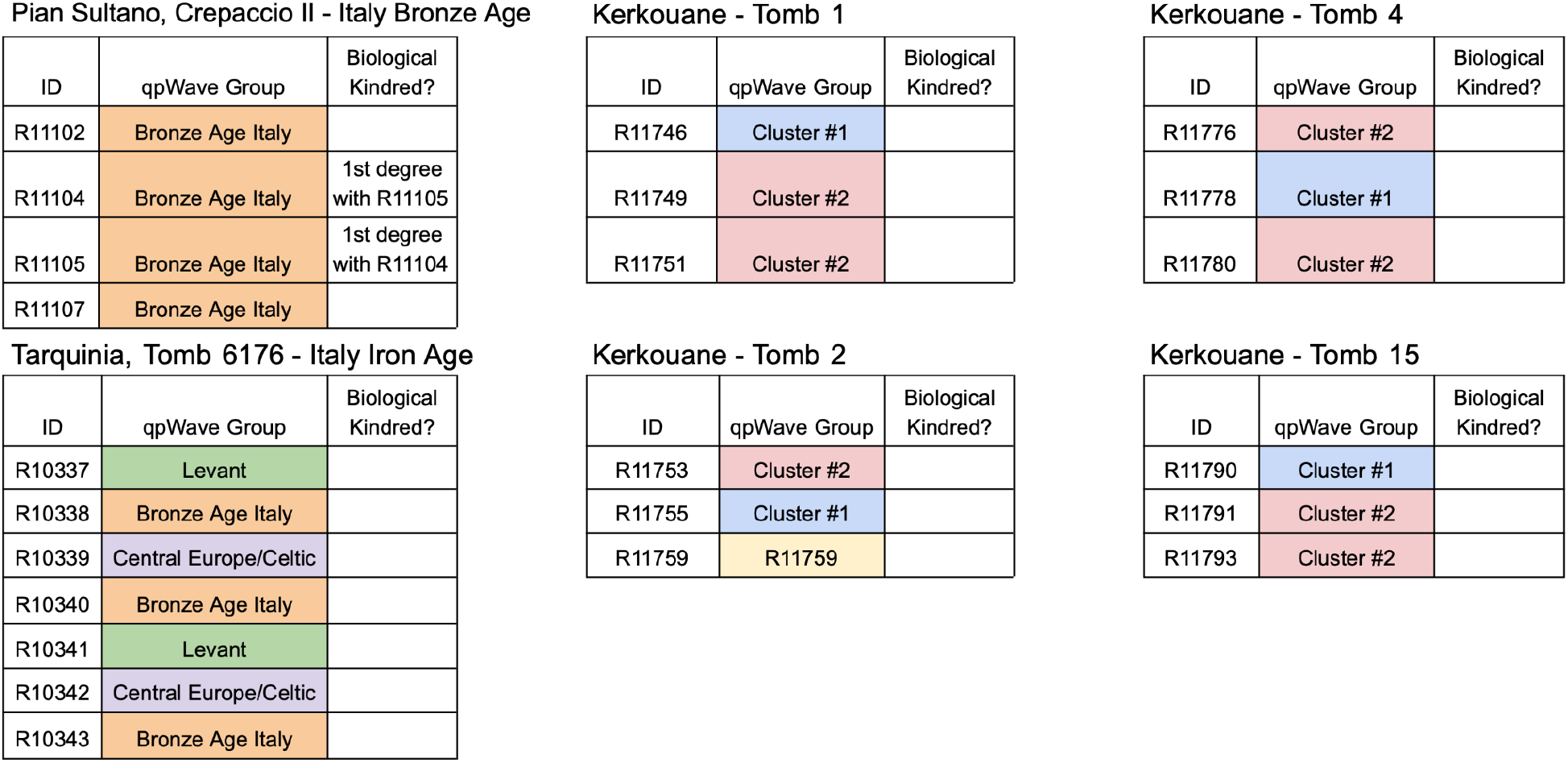
Burial context and biological relatedness information for individuals in this study. Colors show qpWave ancestry groups from Fig. 5.

| ID | qpWave Group | Biological Kindred? |
| --- | --- | --- |
| R11102 | Bronze Age Italy |  |
| R11104 | Bronze Age Italy | 1st degree with R11105 |
| R11105 | Bronze Age Italy | 1st degree with R11104 |
| R11107 | Bronze Age Italy |  |

Tarquinia, Tomb 6176 - Italy Iron Age
| ID | qpWave Group | Biological Kindred? |
| --- | --- | --- |
| R10337 | Levant |  |
| R10338 | Bronze Age Italy |  |
| R10339 | Central Europe/Celtic |  |
| R10340 | Bronze Age Italy |  |
| R10341 | Levant |  |
| R10342 | Central Europe/Celtic |  |
| R10343 | Bronze Age Italy |  |

Kerkouane - Tomb 1
| ID | qpWave Group | Biological Kindred? |
| --- | --- | --- |
| R11746 | Cluster #1 |  |
| R11749 | Cluster #2 |  |
| R11751 | Cluster #2 |  |

Kerkouane - Tomb 2
| ID | qpWave Group | Biological Kindred? |
| --- | --- | --- |
| R11753 | Cluster #2 |  |
| R11755 | Cluster #1 |  |
| R11759 | R11759 |  |

Kerkouane - Tomb 4
| ID | qpWave Group | Biological Kindred? |
| --- | --- | --- |
| R11776 | Cluster #2 |  |
| R11778 | Cluster #1 |  |
| R11780 | Cluster #2 |  |

| ID | qpWave Group | Biological Kindred? |
| --- | --- | --- |
| R11790 | Cluster #1 |  |
| R11791 | Cluster #2 |  |
| R11793 | Cluster #2 |  |

Not only do we see evidence for increasing mobility and interactions across the Mediterranean, we also see possible indications of interactions across the Saharan desert. The sub-Saharan ancestry we observe at Kerkouane may result either from direct contact or indirect contact through the nomadic populations of the Sahara. Trans-Saharan trade routes, made easier by a greener, less arid Sahara than today, had connected the communities of North Africa with their sub-Saharan counterparts since the Bronze Age ^38,39^. Herodotus noted the coexistence of sedentary peoples and nomadic peoples in northern Africa in the 5th century BCE ^40^. In addition to overland networks, these connections to sub-Saharan Africa also occurred by sea. Herodotus described Phoenician trade routes as extending far beyond the Mediterranean to Western Africa via the Atlantic coast and even that a Phoenician and Egyptian expedition had circumnavigated Africa the previous century ^1,2^. The Iron Age may have been a key period for gene flow across the Sahara as well.

Kerkouane was highly cosmopolitan, reflecting the diverse material culture of the city. We observe two groups of individuals who show genetic continuity with the preceding populations of the Maghreb, suggesting standing population structure in coastal North Africa in this period as well as individuals with non-local ancestries. The contribution of autochthonous populations in the region is obscured by the use of terms like “Western Phoenicians”, and even to an extent, “Punic”, in the literature, as it implies a primarily colonial population in North Africa and diminishes local involvement in Iron Age North Africa and the rise of Carthage. As a result, the role of autochthonous populations has been largely overlooked in studies of the Carthaginian world. The high number of individuals with Italian and Greek-like ancestry may be due to the proximity of Kerkouane to Magna Graecia, as well as key trans-Mediterranean sailing routes passing by Cap Bon ^1,41^. Surprisingly, we did not detect individuals with large amounts of Levantine ancestry at Kerkouane. Given the connection of Carthage and its territories as Phoenician colonies, we had anticipated we would see individuals with ancestries similar to Phoenician individuals, such as those published in ^13^. One possible explanation is that the colonial expansion of Phoenician city-states at the start of the Iron Age did not involve large amounts of population mobility and may have been based on trade relationships rather than occupation^42^. Alternatively, this could potentially be due to differential burial practices between local and diasporic communities, such as cremation^1^, or to a disruption in connections between Carthaginian territories and the Eastern Mediterranean, after the annexation of the Phoenician city-states into the Achaemenid Empire.

While Sardinia was comparatively more homogeneous genetically through the Bronze Age than nearby continental regions, such as Iberia and Italy, perhaps due to its insular nature, during the Iron Age, the genomic data from Sardinia shows a rapid increase in heterogeneity in ways that mirror the affiliation of Sardinia as an important part of the Carthaginian Mediterranean, and later of the Roman *mare nostrum*. In the early- and mid-Iron Age, many individual genomes form a clade in pairwise qpWave with the North African population at Kerkouane, while in the late Iron Age, individuals are similar to individuals from mainland Italy and Sicily, reflecting Sardinia’s incorporation into the Roman Empire.

In contrast, in Italy, the majority of studied individuals cluster genetically with the Bronze Age populations of central Italy, indicating a continuity of populations, consistent with the recent findings of ^9^. Alongside this, we observe an increase in heterogeneity, with nearly 40% of the population best modeled with non-local ancestry. Thus, continuity is paired with gene flow patterned by the historical, geopolitical, and even environmental conditions of the time.

The Iron Age in the Mediterranean was characterized by leaps in the ease of seafaring and, as a result, mobility. We see that these technological changes were accompanied by an increase in gene flow and genetic mobility across the Mediterranean, which shaped the ancestry makeup of the populations on its shores. By examining ancient DNA from four archaeological sites in the central Mediterranean, we observe that this technological shift was accompanied by a parallel increase in genetic heterogeneity, in comparison to preceding populations. We observe large amounts of local continuity, with mobility and accompanying gene flow patterned by historical and environmental factors. While further research is needed to make claims about cultural practices, we see indications of admixture between local and diasporic populations, as well as two instances of consanguinity within the diasporic population at Kerkouane. We also suggest that there is a connection between the trend of increasing local heterogeneity and shifts toward modern Mediterranean population structure, indicating that the genetic impacts of mobility were not isolated to port cities, but extended, at least to some degree, to populations inland from the coasts. In *The Making of the Middle Sea*, Cyprian Broodbank notes, “[w]ithout denying the likelihood of various constellations of social, cultural and other identities, early Mediterranean history instead comprises an ever-shifting kaleidoscope of webs of people and practices changing within and between places.” Ancient DNA adds a new approach to examining these webs in the Central Mediterranean during the Iron Age.

## Materials

Short-form site descriptions (see Supplementary Materials for Extended Descriptions)

### Kerkouane

Kerkouane is an exceptionally well-preserved town located on Tunisia’s Cap Bon Peninsula and provides one of the best-surviving windows into Carthaginian daily life ^43–46^. Originally inhabited from 650 - 250 BCE, the population of Kerkouane is thought to have been around 1,200 with an economy primarily based on the production and export of marine resources from the region, including the production and exportation of garum, salt, and Tyrian purple dye derived from locally harvested *Murex sp*. shells ^47^. Kerkouane was abandoned after the 1st Punic War between Carthage and Rome and was never re-inhabited or rebuilt by the Romans resulting in the excellent preservation of the original architecture of the town.

### Sant’Imbenia

Sant’Imbenia, a port town in northwest Sardinia (present-day Alghero) is thought to have been populated by autochthonous Nuragic and Phoenician and Punic people, and has extensive trade contacts with the Eutrsucan city-states of central Italy. This differs from the Phoenician colonies in the south and west of the island, which were thought to be a primarily colonial population. The discovery of a metal workshop and copper ingots at the site suggest it may have been a major center of ore processing.

### Tarquinia

This Etruscan site was one of the largest Iron Age cities in central Italy. It was inhabited throughout the Iron Age and served as one of the primary trading ports between Etruria and the civilizations of the Mediterranean.

### Pian Sultano

The Bronze Age settlement of Pian Sultano is located in central Italy, near modern-day Cerveteri (the Etruscan town of Caere). The earliest record for settlement at the site dates to 2000 BCE. Archaeological investigations of the site have uncovered artifacts indicating Pian Sultano was a farming community that also drew heavily on marine resources. Many ceramics feature design motifs characteristic of the central Italian Apennine culture. Long-distance trade is also indicated in the material culture of the site by obsidian blades, the material for which would have been procured from one of the 4 central Mediterranean island sources - Lipari, Pantelleria, Sardinia, or Sicily ^48^.

## Methods

### Extraction of DNA in Dedicated Cleanrooms

We cleaned, isolated, and powdered the cochlear portion of the petrous bone in dedicated clean rooms following the protocols described in ^49,50^. Following a 30-minute uracil–DNA–glycosylase (UDG) treatment, double-stranded library preparation followed a modified version of the ^51^ protocol (SI, Methods). Libraries passing screening based on DNA concentration were sequenced on an initial next-Seq screening run. Computational authentication of the presence of endogenous ancient DNA was based on 1) the presence of reads mapping to the human genome (hg19 assembly), 2) on the damage patterns at the terminal ends of reads, and 3) contamination analyses using Schmutzi ^52^.

### Newly generated data merged with published data

In total, 30 individual genomes passed endogenous preservation and quality control thresholds (Fig. 1; Dataset S1). Libraries were sequenced on an Illumina NovaSeq 6000 sequencing platform to generate whole-genome shotgun data, with an average genome-wide coverage of 1.1x (range: 0.61 - 1.9x). For the analyses in the paper, we merged the newly generated data reported here with the Allen Ancient DNA Resource v44 ^53^ using PLINK v1.9059. We also added recently published data from Bronze Age Italy to the reference dataset ^9,12,18^. We performed all subsequent analyses on autosomal data.

### Population genetic analyses

Individual biological sex determination was inferred based on the ratio of reads from sex chromosomes and autosome coverages, and kinship analysis was performed using READ (Relationship Estimation from Ancient DNA) ^54^. We carried out *qpAdm* and *qpWave* analyses using ADMIXTOOLS2 ^55^. For modeling the distal ancestries (Fig. 3), we used Mbuti.DG, Russia_Ust_Ishim.DG, CHG, Russia_EHG, Iberia_ElMiron, Czech_Vestonice1, Russia_MA1_HG.SG, Israel_Natufian, Jordan_PPNB as outgroup populations. All individuals in these analyses and the curation of the group labels for these runs can be found in Dataset S2.

### Data availability

All tools and data needed to reproduce and evaluate the conclusions in this paper are presented in the main text and the Supplementary Materials. Alignment files for the DNA sequences for all newly reported individual genomes will be available at the European Nucleotide Archive (ENA) database under the accession number Project PRJEB49419. (www.ebi.ac.uk/ena/browser/view/PRJEB49419).

### Permits to work with archaeological materials

The material reported in this study represents 4 archaeological sites. We have provided the names of the sites, the time periods of occupation, the exact coordinates of the sites, and information about each in the supplementary information. For each site, we worked with the appropriate governmental authorities and permitting bodies to obtain permissions and permits to work on this material. For Pian Sultano and Tarquinia, both located in Lazio Administrative Region of Italy, we received permissions and permits from Daniela De Angelis, the Director of the National Etruscan Museum of Tarquinia (operated by the Direzione Generale Musei Lazio, which is part of the Italian Government’s Ministry of culture) where all material for both Pian Sultano and Tarquinia are housed on December 22, 2018. We also received permits to visit and conduct research in the collection in March and July of 2019. For the material from Sant’Imbenia, Sardinia, we received permissions from Dr. Gabriella Gasperetti, the Archaeological Superintendant of the province of Sassari e Nuoro (Soprintendenza Archeologia, belle arti e paesaggio per le province di Sassari e Nuoro; Sassari, Italy) who oversees these collections on November 1, 2019. For the material from Kerkouane, Tunisia, we received a permit and signed a partnership agreement with the National Heritage Intitute of the Tunisian government’s Ministry of Culture. These were signed and approved by the Director General, Dr. Faouzi Mahfoud on April 4, 2020.

## Supporting information

Dateset S3 - Radiocarbon Dating and Isotopic Analysis Results

Dataset S1 - Newly reported ancient individuals

Dataset S2 - Ancient Reference Genomes.xlsx

Dataset S2 - Ancient Reference Genomes.xlsx

## Acknowledgments

This project was partially supported by the Stanford Interdisciplinary Graduate Fellowship and grants from the Stanford Archaeology Center, The Europe Center Austria Exchange Program and the Stanford Anthropology Dept (HMM); National Science Foundation Graduate Research Fellowship (MA), the Howard Hughes Medical Institute, the Italian Ministry of Foreign Affairs and International Cooperation, and the Istituto Per l’Oriente CA Nallino, and a Ministro dell’Istruzione, Università e Ricerca (MIUR) project grant via the International Association for the Mediterranean and Oriental Studies. We thank all members of the Pritchard lab and Pinhasi lab for their thoughtful and valuable feedback.

## Author Contributions

RP, JKP, AC and MF **designed research**; RP, SS, VO, EP, OC, KO, LD, HMM and DF **performed and supervised laboratory work**; AC and MF **designed the collection strategy for archaeological material**; MF, AC, ML, FLP, FG, FC, DDA, GG, HMM, MC and SA **assembled skeletal material and provided archaeological background**; HMM, MA, SS, JPS, VO, CW, EP, BZ and ZG **curated and analyzed data with input from** JKP, RP, AC, DF and MC; HMM, JPK RP, JPS, MA, SS, CW, MC and ZG **wrote the paper** with input from all collaborators.

## Footnotes

1 Although Phoenician burial practices were thought to have shifted from cremations to interments in the central and western Mediterranean around 650 BCE ^43^, predating the individuals in the study.

