## Supplementary Information for "A Genetic History of Continuity and Mobility in the Iron Age Central Mediterranean"

**This PDF file includes:**

- Supplementary Text
- Figures S1 to S14
- Table S1
- Legends for Datasets S1 to S3
- SI References

**Other supplementary materials for this manuscript include the following:**

- Datasets S1 to S3

Table of Contents

**Supplementary Text** 4

**Site descriptions** 4
- Kerkouane 4
- Sant'Imbenia 4
- Tarquinia 5
- Pian Sultano 6

**Methods** 7
- Date determination and radiocarbon dating 7
- Isotopic Analyses - δ13C (carbon) and δ15N (nitrogen) 7
- DNA extraction and library preparation 8
- Sequencing read processing and library screening 8
- Sample screening criteria 8
- Calling pseudohaploid genotypes 9
- Newly generated data merged with published data 9
- Inference of mitochondrial DNA haplogroups and contamination 9
- Discussion of Uniparental Haplogroups 9
- Genetic Relative Identification 10
- Calculation of heterozygosity 11
- Runs of homozygosity 11
- Principal Component Analysis 11
- qpAdm and qpWave modeling 11
- ADMIXTURE modeling 12
- The Iron Age and the Genetic Structure of North Africa 12

**Supplementary Figures** 14
- Fig. S1. Timeline of all data from the Iron Age central Mediterranean 14
- Fig. S2. Maghreb ADMIXTURE and PCA time series 15
- Fig. S3. Sardinia PCA time series 16
- Fig. S4. Central Italy ADMIXTURE and PCA time series 17
- Fig. S5. Kerkouane PCA and qpAdm admixture modeling 18
- Fig. S6. Admixture modeling for R11759 (Kerkouane Outlier) 19
- Fig. S7. qpWave heatmap with individual ID labels 20
- Fig. S8. qpWave analysis focused on central Italy 21
- Fig. S9. Conditional heterozygosity 22
- Fig. S10. Dietary isotopes across all sites 23
- Fig. S11. Mitochondrial haplogroups 24
- Fig. S12. Mitochondrial haplogroups in absolute counts 25
- Fig. S13.Y-chromosome haplogroups 26
- Fig. S14. Tomb of the Leopards at Tarquinia 27
- Fig. S15. Overview of historic events in the Iron Age Mediterranean 28
- Fig. S16. Kerkouane site map 29

**Supplementary Tables** 30

Table S1. Runs of homozygosity 30

**Supplementary Datasets** **31**

Dataset S1. Newly reported ancient individuals 31

Dataset S2. Ancient individual genomes used in analyses 31

Dataset S3. AMS dating and isotope analysis results 31

Dataset 4. Admixture and cladistic analysis using qpAdm and qpWave 31

Data availability 31

**Supplementary Text**
**Site descriptions**
**Kerkouane**
Geographical coordinates: 36.9615, 11.0804
Analyzed individuals: R11749, R11751, R11776, R11746, R11790, R11793, R11753, R11755, R11759, R11778, R11780, R11791
Date range of individuals reported in this study: 761 - 250 cal BCE

Kerkouane is an exceptionally well-preserved Punic town located on Tunisia's Cap Bon Peninsula and provides one of the best-surviving windows into Carthaginian daily life [1–4]. Due to the destruction of Carthage and many of its territories at the end of the Punic Wars, as well as the continued use and development or rebuilding of these towns under Roman control, insights into Carthaginian daily life are limited.

Originally inhabited from 650 - 250 BCE, the population of Kerkouane is thought to have been around 1,200 with an economy primarily based on the production and export of marine resources from the region, including the production and exportation of garum, salt, and Tyrian purple dye derived from locally harvested Murex sp. shells [5]. Kerkouane was abandoned after the 1st Punic War between Carthage and Rome and was never re-inhabited or rebuilt by the Romans resulting in the excellent preservation of the original Punic architecture of the town.

**Sant'Imbenia**
Geographical coordinates: 40.5938, 8.2045
Analyzed individuals: R11828, R11829, R11835
Date range of individuals reported in this study: 1115 - 774 cal BCE

The archaeological site of Sant'Imbenia lies in Porto Conte Bay, a natural harbor called Portus Nympharum by the geographer Ptolemy in the second century AD, 10 km to the WNW of Alghero (SS). During the fourteenth century BCE,  a single-tower nuraghe was built at Sant'Imbenia, surrounded by huts within the current coastline, and subsequently surrounded by a bastion (6, 7). The nuraghe was in use until the tenth century BCE and the village evolved towards an "urban" organization at the end of the ninth century BCE, around an open space interpreted as a market square. This development is related to seafaring travels around the Western Mediterranean Sea by Phoenicians, Etruscans and Greeks. The site remained a very active center at least until the seventh century BCE, an era of transformations in the indigenous world, which saw the growth of the Phoenician colonies in SW of Sardinia (8–10).

About 900 m SW of the nuragic village, between the first and third centuries CE was built and inhabited the seaside villa of Sant'Imbenia, a large private dwelling dedicated to holidaying and leisure, covering a total area of over 3500 m2, decorated with precious elements, mosaics and refined stucco work on the walls and ceiling, that testify to an opulent lifestyle, found only in Lazio, in Rome, and in the Campi Flegrei area in Campania. Archaeological stratigraphies show that it was still inhabited at least until the 7th century AD, no longer as a luxurious private residence, but sporadically by small social groups, simple pastoral and farming communities (11, 12).

An area used as a necropolis, dating back to I-III century AD, was identified and partially excavated in the 1960s near the road connecting the hamlet of Santa Maria la Palma to Porto Conte. Some burials reused limestone steles with schematically engraved human faces dated to Punic-roman age (13). Later, in 1990s, an emergency archaeological excavation due to the installation of a pipeline was directed by the Superintendence of Archaeological Heritage for the Provinces of Sassari and Nuoro, with the excavation of other burials, mainly human graves, often within amphorae, and sometimes used for multiple burials.

The samples being analyzed come from a stratigraphy damaged by previous land drainage works, plowing and stone removal carried out during the twentieth century. For these reasons, for example, tombs 1, 2, 3, 4 were upset. The necropolis, extremely poor in dating materials, has been referred to the Roman imperial age; however we cannot exclude the presence of remains from a previous period, possibly in a secondary position, due to the conditions of the excavation. As a result of this uncertainty in the dating, we performed radiocarbon dating on all material studied from this context. The individuals reported here date to the the late Bronze and early Iron Age (and others, reported in Antontio et al. 2022, dated to the Roman period).

**Tarquinia**
Geographical coordinates: 42.2542, 11.7576
Analyzed individuals: R10337, R10338, R10339, R10340, R10341, R10342, R10343, R10344, R10359, R10361, R10363
Date range of individuals reported in this study: 759 cal BCE - 3 cal CE

Tarquinia, for the Etruscans Tarchna, was one of the four major urban centers of southern Etruria. It can be traced back all the way to the very origins of the Etruscan people. Located 72 kilometers north of Rome, near the Tyrrhenian Sea coast, Tarquinia was inhabited throughout the Iron Age and served as one of the primary trading ports between Etruria and the civilizations of the Mediterranean[6]. It’s a happy coincidence that the archaeological record coincides with the mythological tale of the founding of the city. This was in the period defined as “Villanovan”, the birth of the Etruscan people.

The complexity and richness of the finds that have come down to us from the Villanovan period are amazing. This is particularly true of the objects from the burial grounds of Arcatelle, Poggio Selciatello, Selciatello sopra, Le Rose and Villa Bruschi Falgari [7]. Over the years these have been added to by the results from archaeological excavations and field surveys on the actual settlement sites, Civita, pianoro dei Monterozzi and Poggio Cretoncini [8,9].

From the end of the ninth to the early eighth century BCE, Tarquinia can be seen, most of all, as a hub, between southern Etruria, southern Italy as a whole and Sardinia. There are also signs of it having become a leading light both culturally and in the political/cultural world. Here there was a consolidated social stratification, with locally crafted merchandise on offer to a world that offered commercial and cultural links.

The port of Gravisca was founded in this period. It provided an opening to the Greek, North African and broader Mediterranean world. This was when Tarquinia’s first funerary paintings appeared, such as the tomb of the leopards, shown in Fig. S13. This period was an exceptional season of painted tombs that graced Tarquinia from about 530 BCE almost nonstop until the Hellenistic period in the fourth to third century BCE. In total, over 6000 tombs have been identified at Tarquinia’s Monterozzi necropolis. About 200 of these tombs include feature wall tombs depicting a range of scenes from Etruscan feasting, daily life, and mythology.

Tarquinia’s role during the fifth century BCE was probably, in part, dulled by sumptuary laws and an internal socio-political transformation of the community. However, this didn’t prevent it from taking part in the Etruscan dodecapolis, the union of the twelve major urban centers. Tarquinia went to war with Rome on more than one occasion, finally being undone at the battle of Sentino in 295 BCE [10].

**Pian Sultano**
Geographical coordinates: 42.0263, 11.9483
Analyzed individuals: R11102, R11104, R11105, R11107
Date range of individuals reported in this study: 2000 - 1600 BCE

The Bronze Age settlement of Pian Sultano is located in central Italy, near modern-day Cerveteri (the Etruscan town of Caere). The earliest record for settlement at the site dates to 2000 BCE. Archaeological investigations of the site have uncovered artifacts indicating Pian Sultano was a farming community that also drew heavily on marine resources. Many ceramics feature design motifs characteristic of the central Italy Apennine culture. Long-distance trade is also indicated in the material culture of the site by obsidian blades, the material for which would have been procured from one of the 4 central Mediterranean island sources - Lipari, Pantelleria, Sardinia, or Sicily [11].

### Methods

#### Date determination and radiocarbon dating

To determine the chronological dates for the individuals in the study, AMS radiocarbon dating was conducted on 60% of the individuals. This includes all three individuals from Sant'Imbenia, 2 individuals from Pian Sultano, 6 individuals from Kerkaoune, and 7 individuals from Tarquinia. We made sure to send an aliquot from at least 1 individual per tomb so that we could obtain an estimate for the use dates of each tomb.

The aliquots sampled consisted of 1 gram of dense petrous bone. These were sent to the W.M. Keck Carbon Cycle Accelerator Mass Spectrometer Lab at the University of California at Irvine. The results were calibrated using the intCal20 calibration curve using the OxCal interface (https://c14.arch.ox.ac.uk/oxcal/OxCal.html). We report these results in Dataset S3.

For individuals not sent for dating, we report both the maximum date range obtained through radiocarbon dating of individuals from the same tomb as well as the archaeologically estimated dates and we used the widest date range available of the two. We made one exception to this for Tomb 6176 at Tarquinia, for which the radiocarbon dates and archaeologically estimated dates were discordant. While the archaeological date estimates place this tomb in the early Iron Age (1000 - 900 BCE), radiocarbon dating in the last two centuries BCE (160 BCE to 3 CE). For these individuals, we used the range of the radiocarbon dates individuals from the tomb.

**Isotopic Analyses -** $\delta$13C (carbon) and $\delta$15N (nitrogen)

Dietary isotopes ( $\delta$13C (carbon) and $\delta$15N (nitrogen)) are informative of food sources a person consumed during their lifetime. We obtained dietary isotopic data for the same individuals submitted for radiocarbon dating at the W.M. Keck Carbon Cycle Accelerator Mass Spectrometer Lab. We report these results in Dataset S3.

Individuals at Kerkouane had high $\delta$15N nitrogen isotope values, indicative of diets high in meat and marine sources of protein, such as fish and shellfish. Marine resources were an important part of the economy of Kerkouane, where Tyrian purple dye and garum were two of the town's major exports. Given its geographic location on Cap Bon Peninsula, Kerkouane also served as an important stop for ships on long-distance voyages across the Mediterranean. One individual, R11780, has very high $\delta$15N nitrogen value compared to other individuals from the same site. Genetically this individually is part of Cluster #2 (non-local ancestry). One possibility is that these values reflect the diet of someone who spent large parts of their life at sea, consuming marine resources.

This is in line with the results from Afat et al 2021, which analyzes the dietary isotopes from 25 individuals form the same necropolis, Arg El-Ghazouani, showing a reliance on C3 plant, no evidence for a heavy reliance on C4 plants, and a highlight variable range of $\delta$15N nitrogen values, potentially indicating variable practices of consuming marine resources, although these variable values did not correlate with age, biological sex or burial context [12].

The individuals at Sant'Imbenia have moderate meat/marine consumption, and those at Tarquinia have lower levels (Fig S11), although it should be cautioned that this cannot be interpreted as dietary differences across sites without a study of the isotopic values of flora and fauna from each site as well, to serve as a background for comparison.

### DNA extraction and library preparation

We cleaned, isolated, and powdered the cochlear portion of the petrous bone in dedicated clean rooms following the protocols described in [13,14]. Using 50mg of bone powder, DNA was extracted by 18-hour incubation of the powder in a solution of Proteinase-K and EDTA. The DNA was then eluted in 50 µl 10 mM Tris-HCl, 1 mM EDTA, 0.05% Tween-20, pH 8.0 as in [15,16].

Following a 30-minute uracil–DNA–glycosylase (UDG) treatment, double-stranded library preparation followed a modified version of the [17] protocol. Libraries were double-indexed with Accuprime Pfx Supermix. The PCR cycling conditions used were as follows: initial denaturation at 95°C for 5 min followed by 12 cycles of 95°C for 15 seconds, 60°C for 30 seconds, and 68°C for 30 seconds with a final elongation at 68°C for 5 min. After indexing, the libraries are purified using the MinElute system (Qiagen) and eluted in 25uL of 1 mM EDTA, 0.05% Tween-20.

Libraries passing screening based on DNA concentration were sequenced on an initial next-Seq screening run. Computational authentication of the presence of endogenous ancient DNA was based on 1) the presence of reads mapping to the human genome (hg19 assembly), 2) on the damage patterns at the terminal ends of reads, and 3) contamination analyses using Schmutzi [18], as done in [19].

We created and screened libraries from 23 Estrucan individuals (16 from the site of Tarquinia, 4 from Nepi Sante Grotte Gigliastro and 4 from Vulci Osteria), with 11 libraries (all from Tarquinia) passing the endogenous preservation and quality control measures described above. From Kerkouane, we created libraries for 20 individuals, with 12 successfully passing preservation and quality control standards. All libraries created for Sant'Imbenia (n=3) and Pain Sultano (n=4) successfully passed our screening thresholds.

### Sequencing read processing and library screening

All libraries were initially sequenced to low coverage using Illumina NovaSeq SP in order to screen for endogenous DNA preservation and authenticity (details of criteria in the following section). Libraries that passed screening criteria were sequenced on an Illumina NovaSeq 6000 to generate whole-genome shotgun data. Sequencing data was processed and pseudohaploid genotypes were called as done in [19], and summarized as follows. Following demultiplexing, adapters were removed, reads were filtered (Mapping Quality > 30, Minimum Length > 30), and the two base pairs on each end of the reads were trimmed using Cutadapt (v1.14) [20]. Reads were then aligned to hg19 using bwa (0.7.15-r1140) [21]. For each library, aligned reads were sorted by coordinate using Picard's SortSam (version 2.9.0-1-gf5b9f50-SNAPSHOT) and read groups were added using Picard's AddOrReplaceReadGroups (version 2.9.0-1-gf5b9f50-SNAPSHOT) (http://broadinstitute.github.io/picard/). Aligned read data from different libraries for the same sample were merged and then deduplicated using samtools rmdup (http://www.htslib.org/doc/samtools.html) [22]Genome-wide and chromosomal coverage were assessed using depth-cover (version 1.0.3, https://github.com/jalvz/depth-cover). Samples had an average genome-wide coverage of 1.1x (range: 0.61 - 1.9x)

### Sample screening criteria

Samples were screened and selected using the following criteria: 1) >10% reads aligned to the hg19 build of the human genome; 2) a C>T mismatch rate at the 5'-end and G>A at the 3'-end of the sequencing read of 5% or above (characterized with mapDamage v2.0.8) [23]; 3) with a contamination level <= 3%. The C>T and G>A mismatch rates were assessed using the ends of reads prior to trimming two base pairs (as described in the previous section). Since samples only underwent partial-UDG treatment, the ends of the reads still contain the molecular damage signature characteristic of ancient DNA. Contamination rates were estimated with three methods: 1) damage pattern and polymorphism in mitochondrial DNA with Schmutzi [18], 2) atypical ratios of coverages of X and Y chromosomes to autosomes calculated with ANGSD [24] and 3) for male samples, high heterozygosity on non-pseudo-autosomal region of the X chromosome with the "contamination" tool in ANGSD [24]. Individual biological sex determination was inferred based on the ratio of reads from sex chromosomes and autosome coverages [25].

**Calling pseudohaploid genotypes**

Pseudohaploid genotypes for individuals in this study were called using the pipeline and tool created by Stephan Schiffels (https://github.com/stschiff/sequenceTools). Samtools mpileup was used to generate read coverage of a select SNP. For maximum overlap with published ancient and modern samples, variants were selected based on those used in the Human Origins panel and in the modern reference panel published by Lazaridis et. al. 2014 [26]. This resulted in a total of 481,259 SNPs. A filter of minimum base and mapping quality of 30 (--min-BQ and --min-MQ) were also applied during samtools mpileup. Pseudohaploid genotypes were called by randomly choosing one allele from each site where there was read coverage, using pileupCaller.

**Newly generated data merged with published data**

In total, 30 individual genomes passed endogenous preservation and quality control thresholds (Fig. 2; Dataset S1). For the analyses in the paper, we merged the newly generated data reported here with the Allen Ancient DNA Resource v44 (49) using PLINK v1.9059. We also added recently published data from Bronze Age Italy to the reference dataset (17). We performed all subsequent analyses on autosomal data.

**Inference of mitochondrial DNA haplogroups and contamination**

Schmutzi [18] was used to estimate mitochondrial haplogroups and mitochondrial-based contamination estimates, as in [19]. For determining mitochondrial contamination and the endogenous mitochondrial genome, we used untrimmed reads to identify and extract only reads with a damage signature. The endogenous consensus mitochondrial genome was called simultaneously while estimating mitochondrial contamination using schmutzi [18]. Sequencing reads were aligned to the revised Cambridge Reference Sequence (rCRS) mitochondrial genome (NC_012920.1).

We used the contDeam tool in schmutzi with the following parameters: length of expected deamination set to 2 (--lengthDeam 2) and library type set to double strand (--library double). Schmutzi was used to 1) estimate contamination based on a haplogroup frequency database in tandem with the deamination estimates from contdeam and 2) to assemble the endogenous consensus mitochondrial genome informed by contamination. Base quality filtering of 30 (--qual 30) and the --uselength parameter were both used. This provided a contamination estimate based on deamination rates, a contamination estimate based on haplogroup frequencies, a contaminant mitochondrial genome, and an endogenous mitochondrial genome.

Haplogroups for the endogenous mitochondrial genomes were called using the command line version of Haplogrep (v2.1.20) [27]. Contamination estimates are reported along with X chromosome contamination estimates, where possible (for males). Endogenous haplogroups and contamination estimates are reported in Dataset S1. Endogenous haplogroups are visualized in Fig. S11 and S12.

**Discussion of Uniparental Haplogroups**

<u>Kerkouane</u>

At Kerkouane, of the 7 individuals, 4 female and 3 male, in the non-local cluster (cluster #2), 6 individuals carry either HV or H. Haplogroup HV is thought to have originated in Mediterranean European during the last glacial maximum (LGM), and haplogroup H is thought to have originated in southeastern Europe at a similar time period (citation). This haplogroup has not been reported in earlier populations from the Maghreb, although existing data is limited. The seventh individual carries mitochondrial haplogroup I5, which is thought to have originated in western Asia. Of the 4 individuals in cluster #1, two have haplogroups indicative of continuity with preceding Maghrebi populations, supporting the observations of continuity from the autosomal analyses. R11778 carries haplogroup L3e2bA, which is most common in western Africa, indicating potential trans-Saharan connections already existing in this period [28]. Finally, R11790 carries HV13 which, like the individuals in cluster #2 carrying HV which implies admixture between local and a non-local female carrying this haplogroup. R11759, the outlier individual carries, U6, a common northern African haplogroup, supports standing population structure.

Of the 12 individuals at Kerkouane, 4 individuals are male. Three of these individuals, all from cluster #2 (non-local) all carry J2b, that has previously been associated with the Phoenician expansion [29]. This differs from what is seen in the nuclear and mitochondrial analyses. This could be a two-step process, of this haplogroups dispersing in the Mediterranean and then arriving at Kerkouane. Alternatively, J2b is also found in the BA/Nuragic Sardinian individuals and may reflect the close ties between the Maghreb and Sardinia. Finally, one individual from cluster #1 carries haplogroup R1b, which is associated with Bell Beaker cultures in Europe and common in central Italy at this time and became prevalent in southern Iberia by the Bronze Age (Villalba-Mouco et al., 2021). This supports the possibility that the steppe component observed at Kerkouane may have been introduced through mobility occurring in the central and western Mediterranean, as also suggested by Fernandes et al. 2020.

Sant'Imbenia

All three mitochondrial haplogroups observed, HV0a, K1a2, and U5b3a2 have been reported in for Sardinian individuals from earlier periods, supporting continuity at between these late Bronze Age/early Iron Age individuals and the preceding population of the island. Of the two Y-chromosome haplogroups, I2 is common in Neolithic and Chalcolithic Iberia, France, Hungary, and H3 is a relatively uncommon in ancient individuals, but has been reported in Neolithic Anatolia and early Bronze Age France.

Tarquinia

In central Italy, we observe a dramatic increase in the Y-haplogroup R1b, which has been identified in the Bell Beaker cultures of Europe. R10341 carries Y-chromosome haplogroup J1a, which is found most commonly among Bronze and Iron Age Anatolian and Levantine populations. This haplogroups has not been observed in preceding periods in Italy. This individual has predominantly Levantine autosomal ancestry. R10363 carries G2a, which was also present in Bronze Age individuals from central and northern Italy [30]

**Genetic Relative Identification**

Genetic relative identification was performed using READ (Relationship Estimation from Ancient DNA) [31]. We identified one pair of related individuals, R11104 and R11105, consistent with a first-degree relationship (i.e. parent and child, or two siblings). These two individuals were buried together in tomb 2 at the Bronze Age site of Pian Sultano, along with two other unrelated individuals. Genomic data for all 4 are reported in this study. Interestingly, although a number of individuals from the Iron Age sites (Table 1) were also interred in shared burials, none of these individuals were determined to be first- or second-degree biological kin.

**Calculation of heterozygosity**

We calculated heterozygosity using variants that are already known to be segregating in human populations, following the same approach as in [19]. Three study samples, all from Iron Age Kerkouane

(Tunisia), had heterozygosity beyond one standard deviation of the population average for that region. R11759 has high heterozygosity. R11753 and R11791, have low heterozygosity (Fig. S9).

**Runs of homozygosity**
We followed the same approach as in [19] to estimate recent inbreeding by calculating runs of homozygosity (ROH). Only four samples had more than one ROH segment of 5Mb or longer (Table S2). Two individuals from Kerkouane, R11753 and R11791, show strong evidence of inbreeding, with ROH segments over 50Mb in length. In both cases, with over 12% of the genome being homozygous (segments > 5Mb in length), the parents were likely 2nd-degree relatives. Consistent with the homozygosity analysis, both individuals also have very low conditional heterozygosity (Fig. S9). While it is known that endogamous marriage practices were common in the contemporaneous ancient Greek world, less is known about such practices in the Carthaginian world, due to fewer surviving written records [32]. The other two individuals with more than one ROH segment (R11778 and R10339) have little evidence of inbreeding as only less than 1% of their genome is homozygous.

**Principal Component Analysis**
We generated a Principal Component Analysis (PCA) reference space based on modern populations from around the Mediterranean, Europe, the Middle East, and North Africa using the tool smartpca from the EIGENSOFT package version 8.0.0 [33]. The list of these modern populations and corresponding individual IDs can be found in Dataset S2. We project a set of ancient individuals, primarily from the Bronze and Iron Age Mediterranean, onto the modern reference space (Fig. 3, Fig. 4A). We used the parameter “numoutlieriter:0” to retain all outlier individuals in the projections, and “shrinkmode: YES” to correct for shrinkage towards the origin when estimating  PC scores.

**qpAdm and qpWave modeling**
We carried out *qpAdm* and *qpWave* analyses using ADMIXTOOLS2 [34]. ADMIXTOOLS2 uses F2 statistics as the basis for all downstream analysis. For qpWave, we used pre-computed F2 statistics, computed with the maxmiss parameter set to 0.5 on a set of individuals with greater than 0.5x coverage (to reduce the number of individuals with a low number of shared snps). For both qpWave pairwise analyses (Supplementary Table 4, Tab3), we used the default settings of ADMIXTOOLS2. ADMIXTOOLS2 automatically differentiates between diploid and pseudohaploid data. Since we are using pseudohaploid data, the program functions similar to the Admixtools, Inbreed=YES option.

For modeling the distal ancestries using qpAdm (shown in Fig. 4, S5), we used Mbuti.DG, Russia_Ust_Ishim.DG, CHG, Russia_EHG, Iberia_ElMiron, Czech_Vestonice1, Russia_MA1_HG.SG, Israel_Natufian, Jordan_PPNB as outgroup populations, referred to here as “Right Set A”. The IDs for each individual in each of these outgroup populations can be found in Dataset S2. For both qpAdm modeling, we ran ADMIXTOOLS2 with the allsnps = TRUE setting and the Inbreed=YES option.

We chose a set of distal source populations previously shown to be informative for understanding the diversity of the Mediterranean during this period for qpAdm admixture modeling, including Western Hunter-Gatherer (WHG), Yamnaya Samara, Anatolian Neolithic, Iranian Neolithic, and Late Neolithic farmers from Morocco, referred to here as “Left Set A” [19,35,36]. To find the best fitting combinations of distal sources, we use qpadm_rotate() to compare all possible combinations of these source populations for each individual. In each iteration of the rotation, any of the potential source populations from Left Set A not used as sources in the model are added to the list of outgroup (right) populations. We then selected all models with a p-value greater than 0.05 with the fewest number of source populations. This could potentially be more than one model, and we indicated in figures and the text when this is the case. We then ran qpAdm() on each working model to calculate the ancestry proportions, as well as standard errors.

For R11759, there were no working distal qpAdm models with the original set (Left Set A) of 5 distal source populations (Fig. 5). We replaced Morocco Late Neolithic with Morocco Early Neolithic farmers and a hunter-gatherer individual from Ethiopia from ~4500 BP, both of which produced working models. Using competition modeling (where possible sources are rotated to the outgroup), the best model uses ~70% Morocco Early Neolithic ancestry and ~30% Anatolia Neolithic (Fig. 4, Fig. S6), and models with the hunter-gatherer individual from Ethiopia from ~4500 BP (from Mota Cave), no longer pass the significance threshold of p-value > 0.05 when other populations from sub-Saharan Africa are added to the outgroup list. For better resolution on the potential sub-Saharan African ancestry source population for R11759, we have added additional populations, specifically, Kenya_LukenyaHill_3500BP, Kenya_MoloCave_1500BP, Kenya_PastoralN, to the models, both as a potential source populations and to the outgroup list. These additions did not produce any additional working models as sources, but "break" (i.e. result in models with p-values below the 0.05 threshold) the models with Morocco Early Neolithic and Ethiopia from ~4500 BP, suggesting that gene flow between these populations may be violating the assumed tree structure of qpAdm models.

We used the union of Right Set A and Left Set A as reference populations to perform pairwise qpWave on a set of Bronze and Iron Age individuals to test whether each individual in the pair can be modeled with the same ancestry components in qpAdm in comparison to a set of reference populations. Clustering was generated using 1 - log(p-value) to calculate distances between each individual, with clusters called by the cutree tool in R. This approach tests whether each pair of individuals can be modeled using the same ancestry components in qpAdm. This helps us identify groups of individuals from different geographic locations who can be modeled similarly. Given the genetic heterogeneity that characterizes the Iron Age Mediterranean, these groupings identified in qpWave help identify genetically similar individuals across geographic regions.

**ADMIXTURE modeling**

We used supervised ADMIXTURE modeling to interpret the study individuals as a mixture of source populations [37]. To determine the appropriate populations and number of populations to use as "sources" for supervised admixture, we performed unsupervised admixture for k = 2 through 7, with 5-fold cross validation, and 4 repetitions of each k. Runs for k = 3 through 5 had the lowest cross validation errors across repetitions. As a result of these runs, we selected 5 source populations that each maximized a unique component at k = 5 in the unsupervised run: Western Hunter-Gatherers (WHG), Anatolia Neolithic Farmers, Iranian Neolithic Farmers, Moroccan Hunter-Gatherer and Early Neolithic Farmers, Eneolithic Steppe Herders. A list of all individuals used to represent each population in the analyses can be found in Dataset S2.

**The Iron Age and the Genetic Structure of North Africa**

The Iron Age appears to be a key period for the formation of the current genetic structure of North Africa. Previous research suggests present-day central and western North African populations can be modeled as having four primary ancestry components: a local/autochthonous Maghrebi component derived from paleolithic hunter-gatherer populations in the region[38–40]; a Near Eastern component thought to have been introduced with Arab rule of the region in the Medieval period[41]; a sub-Saharan African component [38,42]; and a European component. While many papers have suggested the Near Eastern and European components resulted from recent historical movements, such as Arab rule in Medieval North Africa and trans-Mediterranean trade in the last 500 years, Fregel et al. 2018 has shows the "European" component is, at least partially, linked to the farming expansion and is similar to Anatolian and early European farmers[38]. We show that both Near Eastern and sub-Saharan African components were present in North Africa earlier than previously thought, reflecting the ongoing interconnectedness of North Africa to these regions for millennia [38,43–45], which supports the findings from research on the heterogeneity of mitochondrial haplogroups in the Canary Islands, showing the presence of haplogroups characteristic of sub-Saharan Africa[46].

**Supplementary Figures**

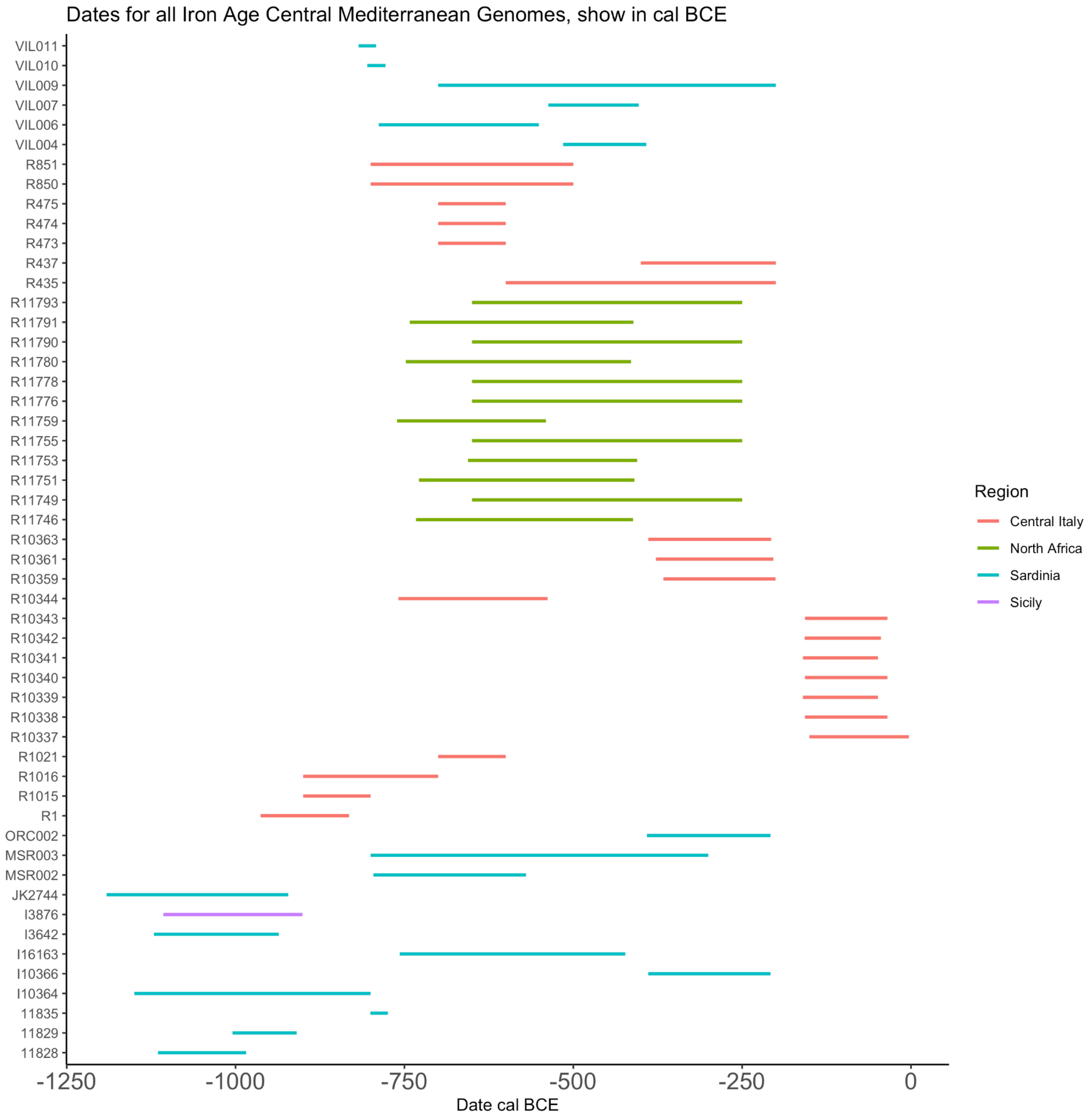


**Fig. S1. Timeline of all data from the Iron Age central Mediterranean**
This includes new data, plus data from [19,36,47]. Dates shown are based on radiocarbon dates when available, and otherwise on archaeological date estimates (see Datasets S1, S2, S3).

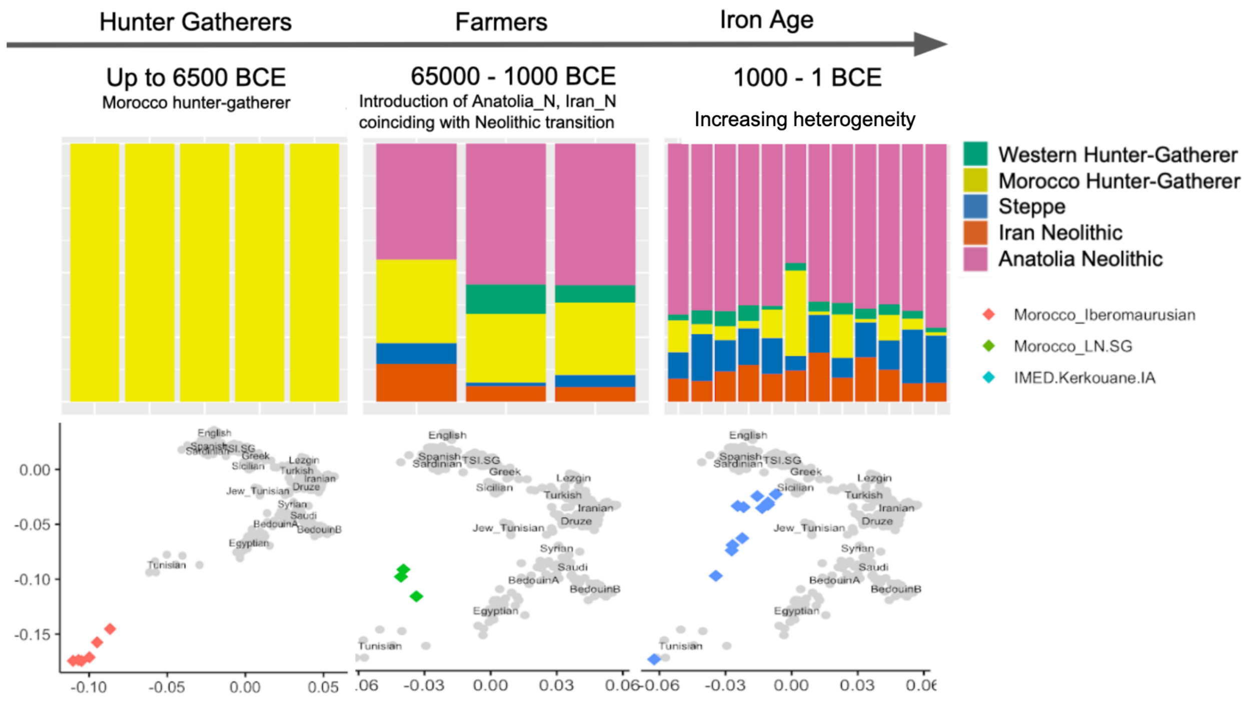


**Fig. S2. Maghreb ADMIXTURE and PCA time series**
A time series of new and published ancient genomes from North Africa, analyzed using ADMIXTURE and PCA. All new genomes are shown in the Iron Age column. For a larger PCA, see Fig. 3.

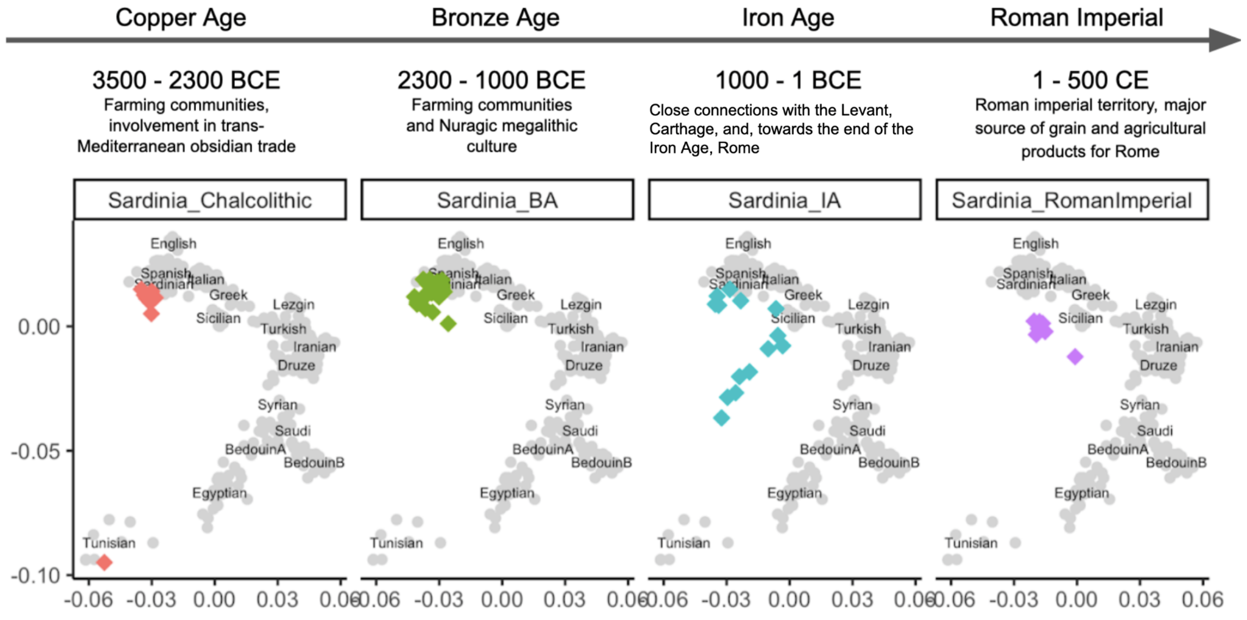


**Fig. S3. Sardinia PCA time series**

A time series of new and published ancient genomes from Sardinia, analyzed using PCA. This plot shows relatively homogeneous ancestry in the Chalcolithic and Bronze Age, followed by a shift toward North African ancestry in the Iron Age (when the island was part of the Carthaginian Empire) and towards mainland Italy, Sicily and Greece in the Imperial Period (when Sardinia was part of the Roman Empire). These shifts seem to mirror the geopolitical affiliations of Sardinia. All new genomes are shown in the Iron Age column. For a larger PCA, see Fig. 3.

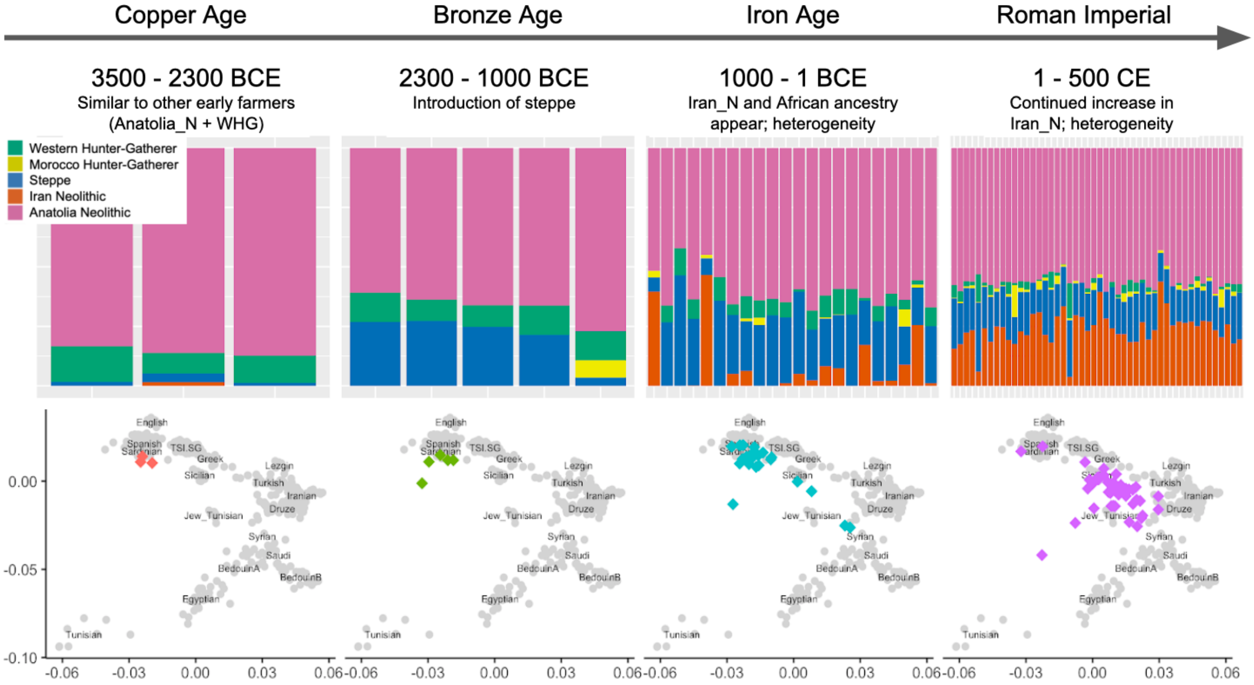


**Fig. S4. Central Italy ADMIXTURE and PCA time series**

Unsupervised ADMIXTURE plot of genetic data from central Italian individuals organized chronologically.

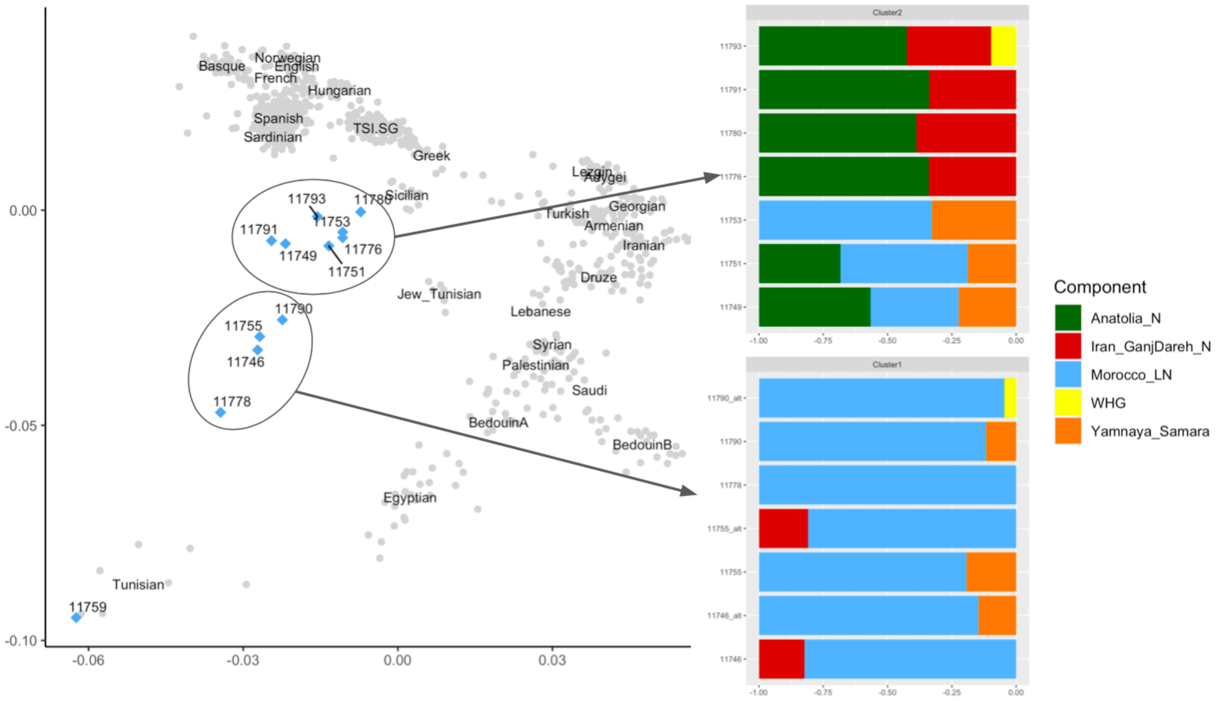


**Fig. S5. Kerkouane PCA and qpAdm admixture modeling**
Kerkouane PCA and qpAdm admixture modeling.

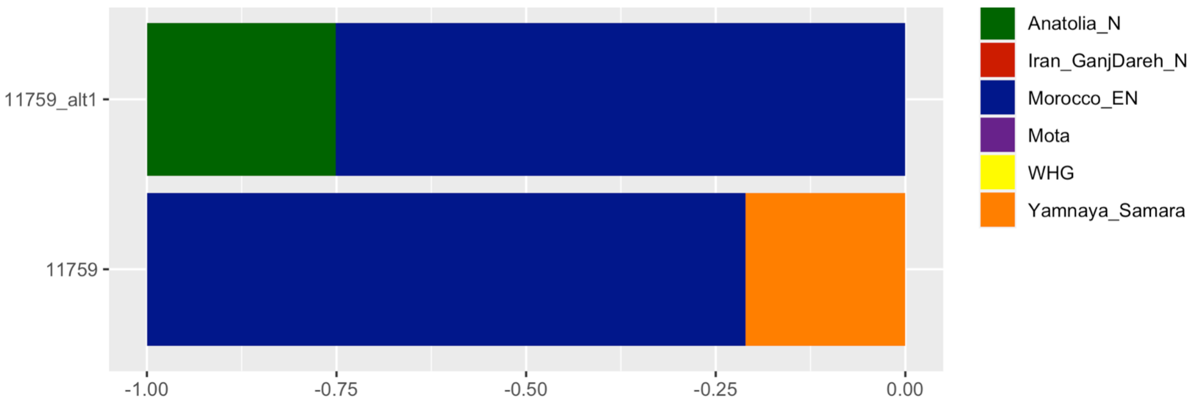


**Fig. S6. Admixture modeling for R11759 (Kerkouane Outlier)**
11759 is the outlier individual that projects in PCA onto modern Tunisian individuals. There are no working models for this individual with either Morocco_LN or Morocco_Iberomaurusian, but two possible models including Morocco_EN as sources produce working models, both of which are shown above.

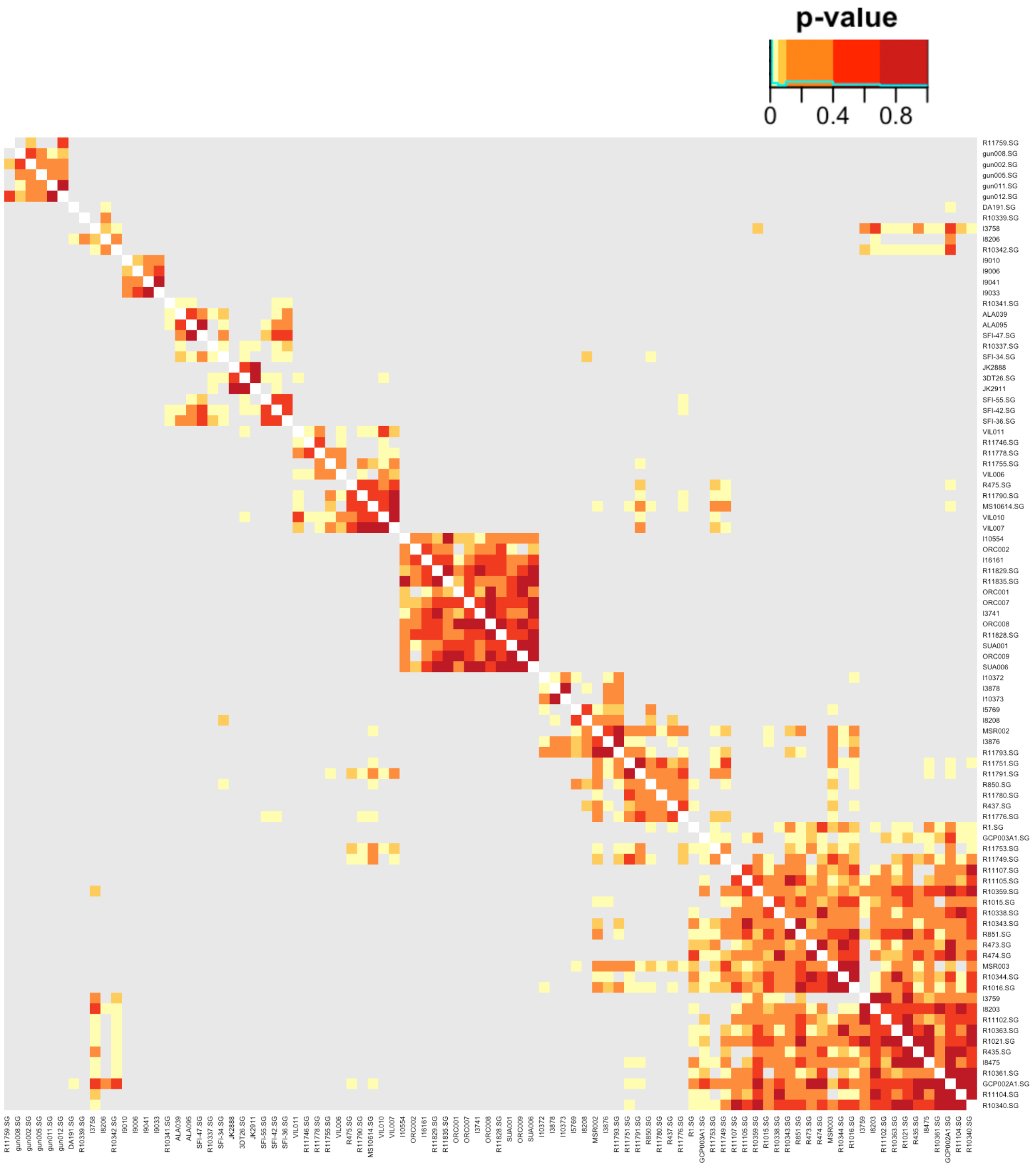


**Fig. S7. qpWave heatmap with individual ID labels**

qpWave heatmap shown in Fig 5, with individual IDs included as labels. The plot shows the clustering of Bronze Age and Iron Age genomes from the central Mediterranean with other relevant ancient individuals. Heatmap colors represent the p-values for each pair-wise model. Values over 0.01 (shown in yellow) indicate that a pair of individuals can be modeled with the same ancestry components in qpAdm in comparison to a set of reference populations ( Mbuti.DG, Russia_Ust_Ishim.DG, CHG, Russia_EHG, Iberia_ElMiron, Czech_Vestonice1, Russia_MA1_HG.SG, Israel_Natufian, Jordan_PPNB, Western Hunter-Gatherer (WHG), Yamnaya Samara, Anatolian Neolithic, Iranian Neolithic, and Late Neolithic farmers from Morocco).

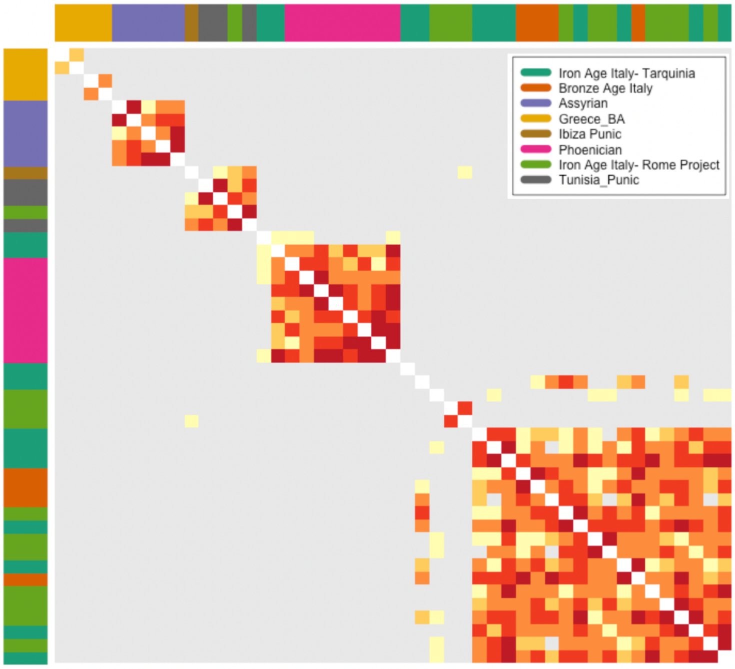


**Fig. S8. qpWave analysis focused on central Italy**
Localized qpWave analysis, focusing on the Italian Bronze and Iron Age individuals.

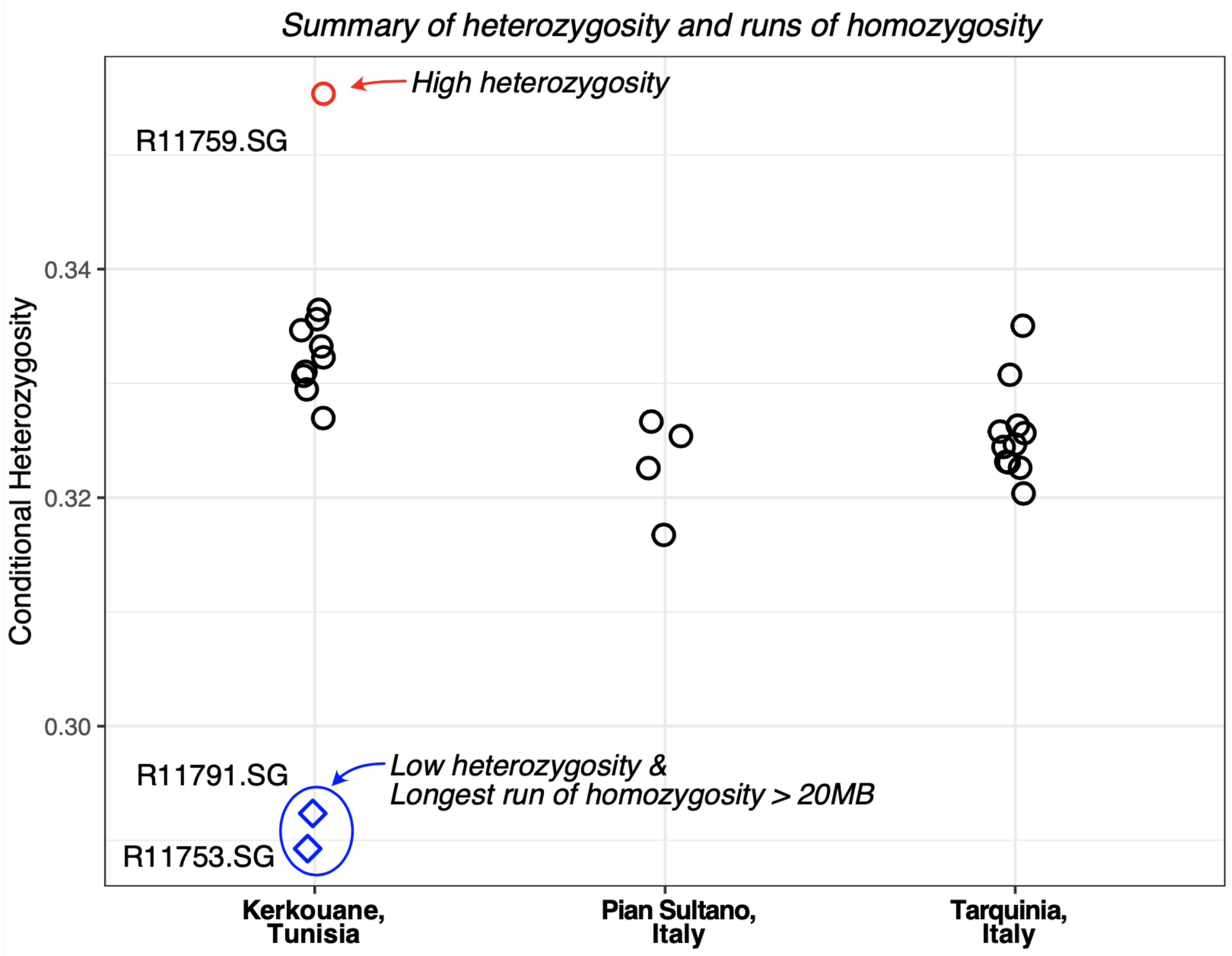


**Fig. S9. Conditional heterozygosity**

We calculated heterozygosity using variants that are already known to be segregating in human populations, following the same approach as in [19].

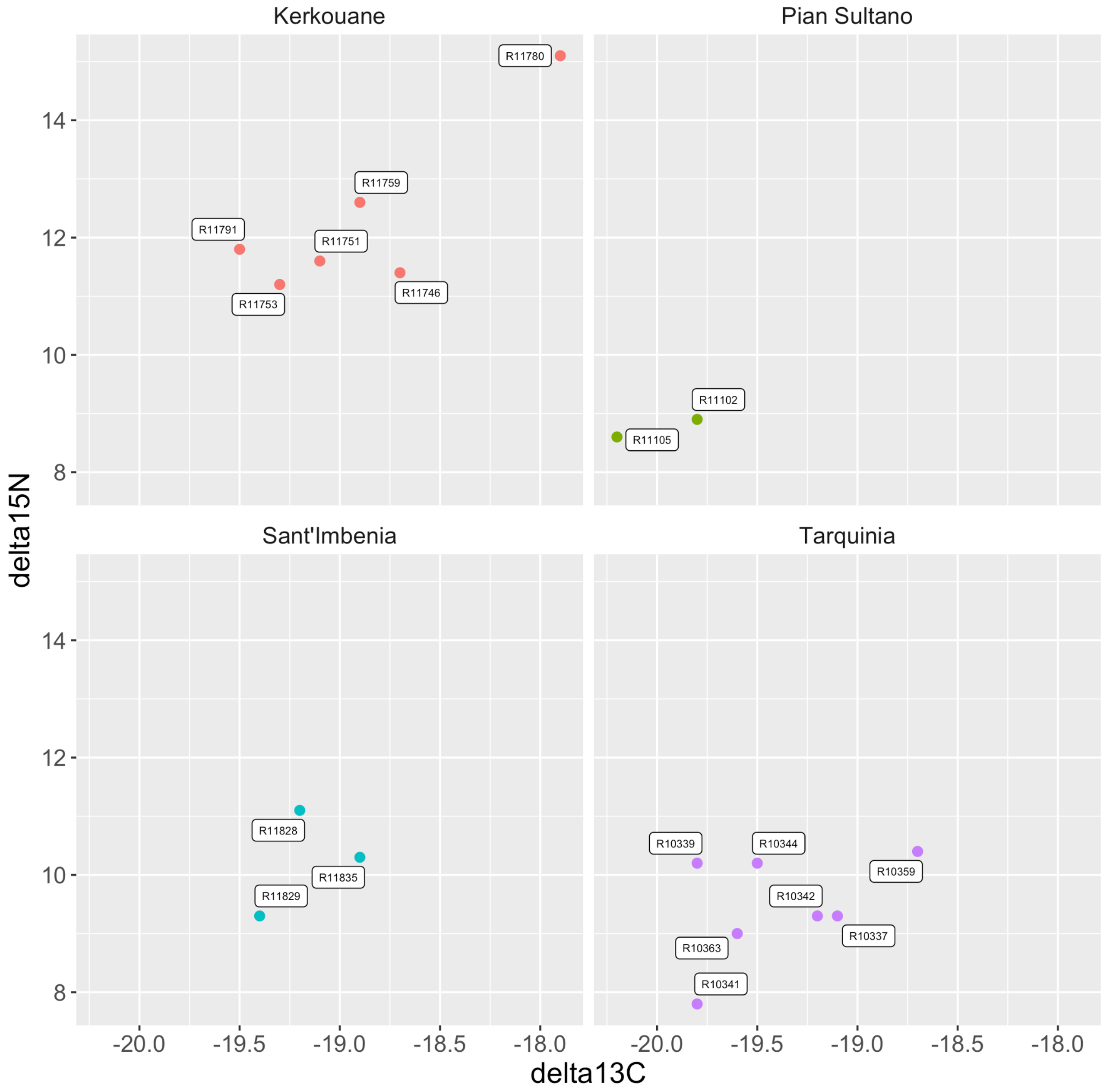


**Fig. S10. Dietary isotopes across all sites**
Dietary Isotopes for individuals across all sites studied.

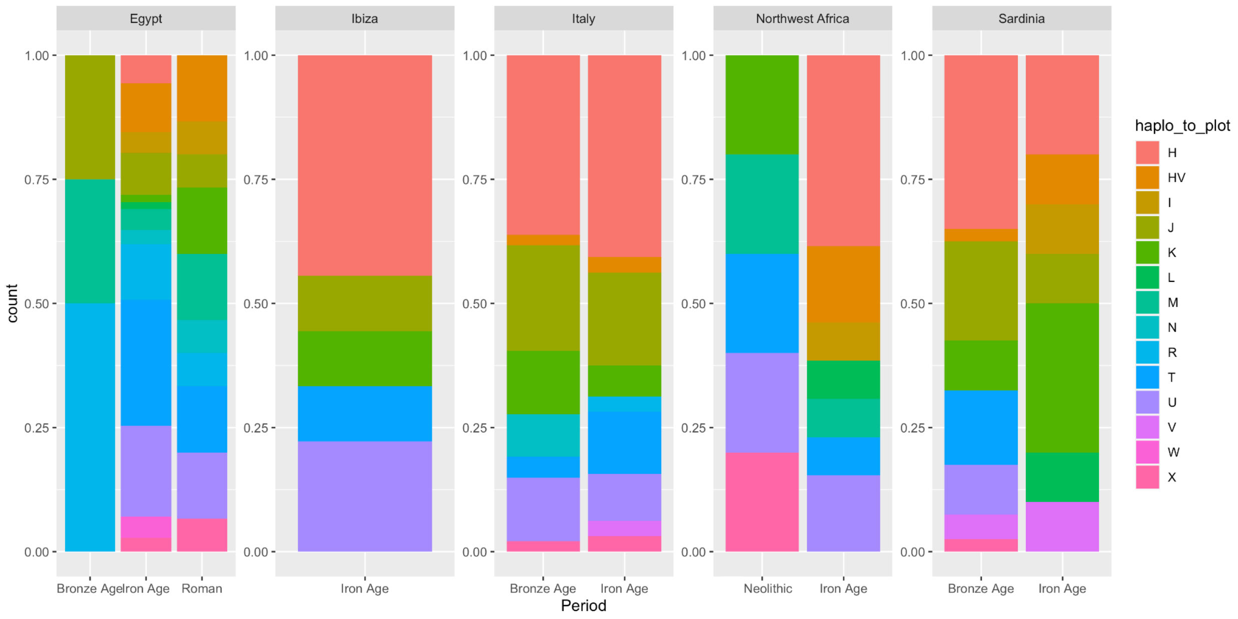


**Fig. S11. Mitochondrial haplogroups**

Mitochondrial haplogroups for each region in the Iron Age Central Mediterranean and the preceding period, show in percent of population.

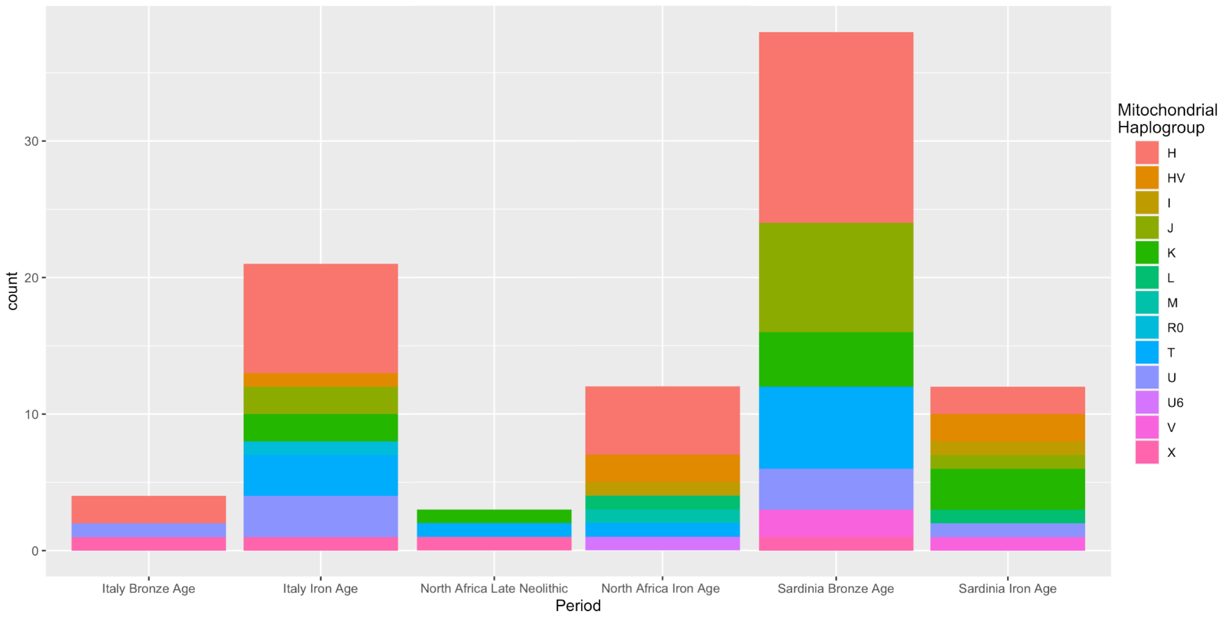


**Fig. S12. Mitochondrial haplogroups in absolute counts**
Mitochondrial haplogroups for each region in the Iron Age Central Mediterranean and the preceding period, shown in absolute counts.

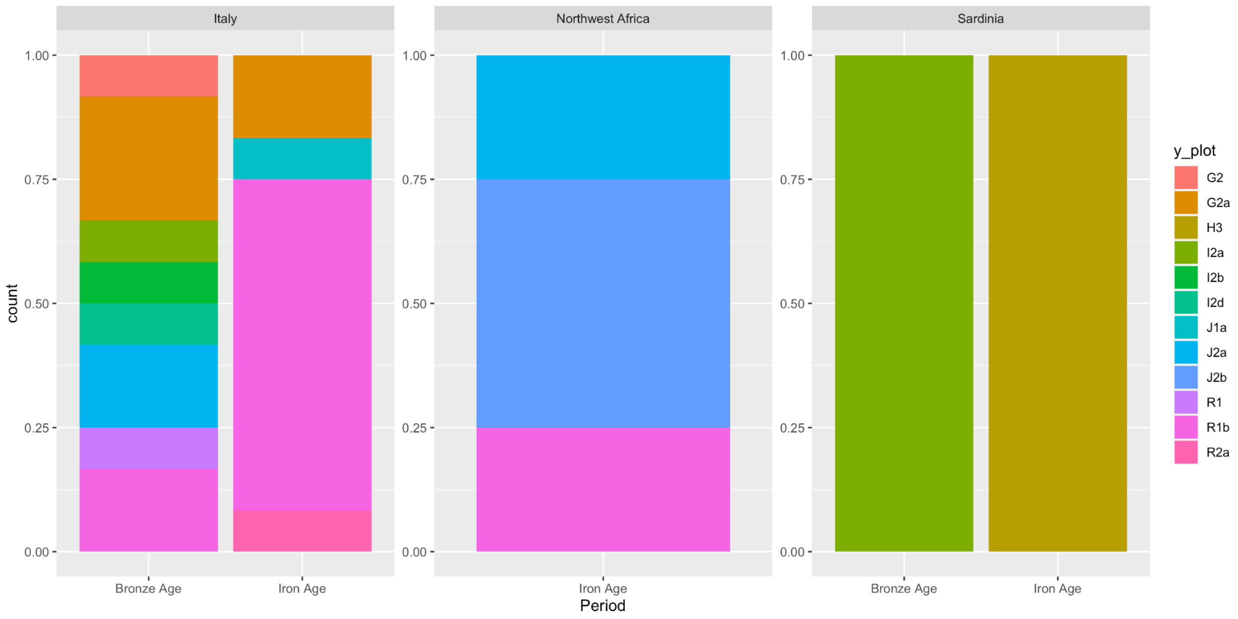


**Fig. S13.Y-chromosome haplogroups**

Haplogroups for all newly reported individuals, shown for each region in the Iron Age Central Mediterranean and the preceding period, show in percent of population.

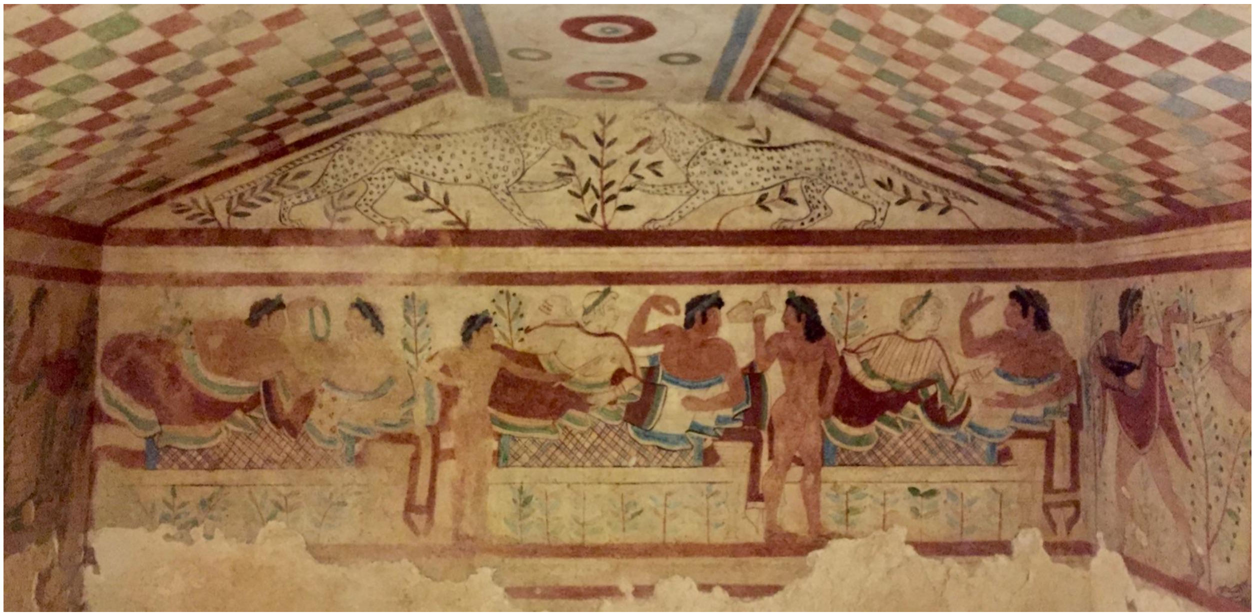

**Fig. S14. Tomb of the Leopards at Tarquinia**

One of the painted tombs at Tarquinia, depicting banqueters feasting. This fresco dates to the 5th century BCE. Photograph taken by H. Moots.

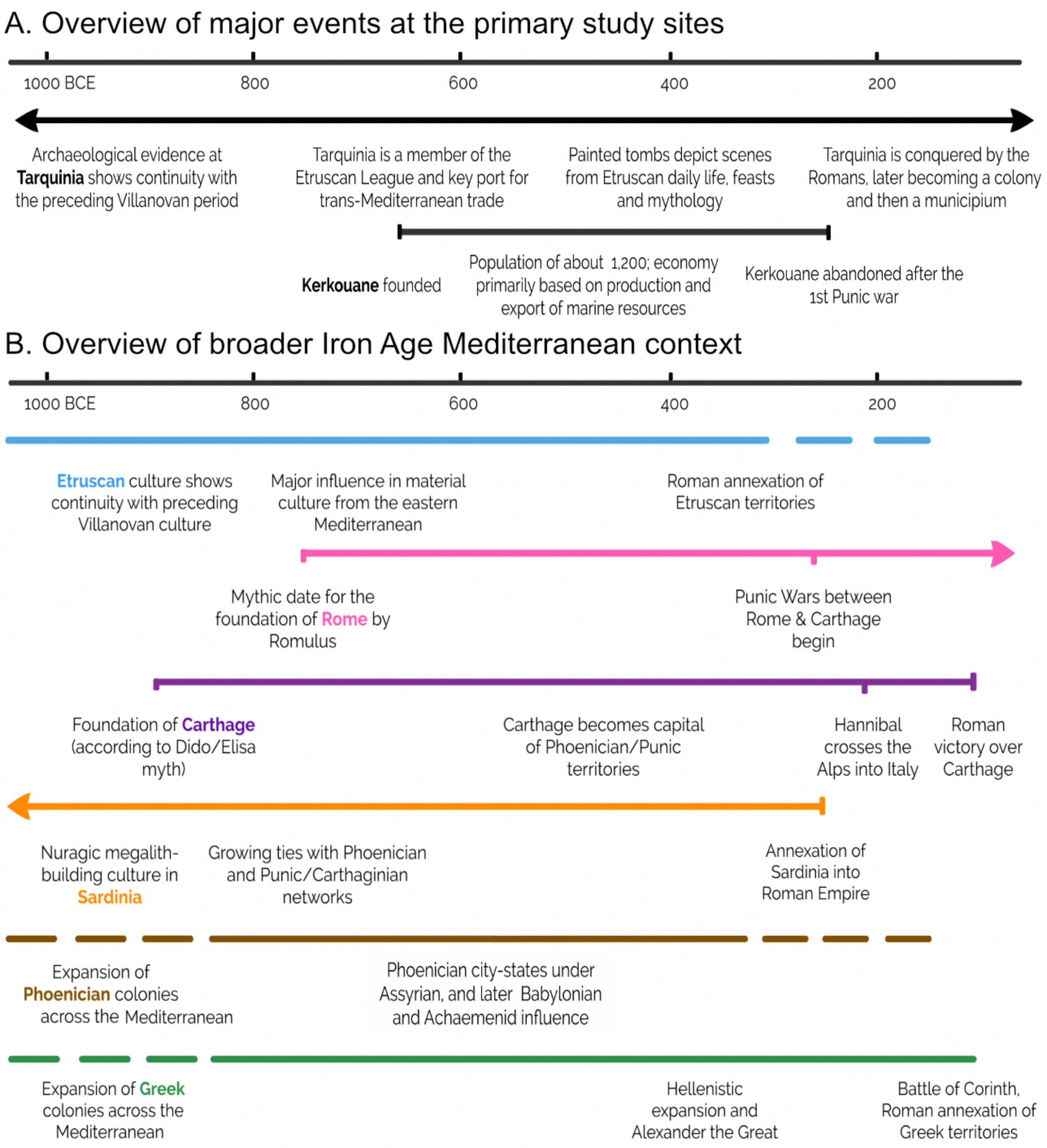


**Fig. S15.** **Overview of historic events in the Iron Age Mediterranean**

Each line corresponds to a cultural group, indicated in the labels below the lines. Dashed lines represent gradual processes that do not have a definitive start or end date.

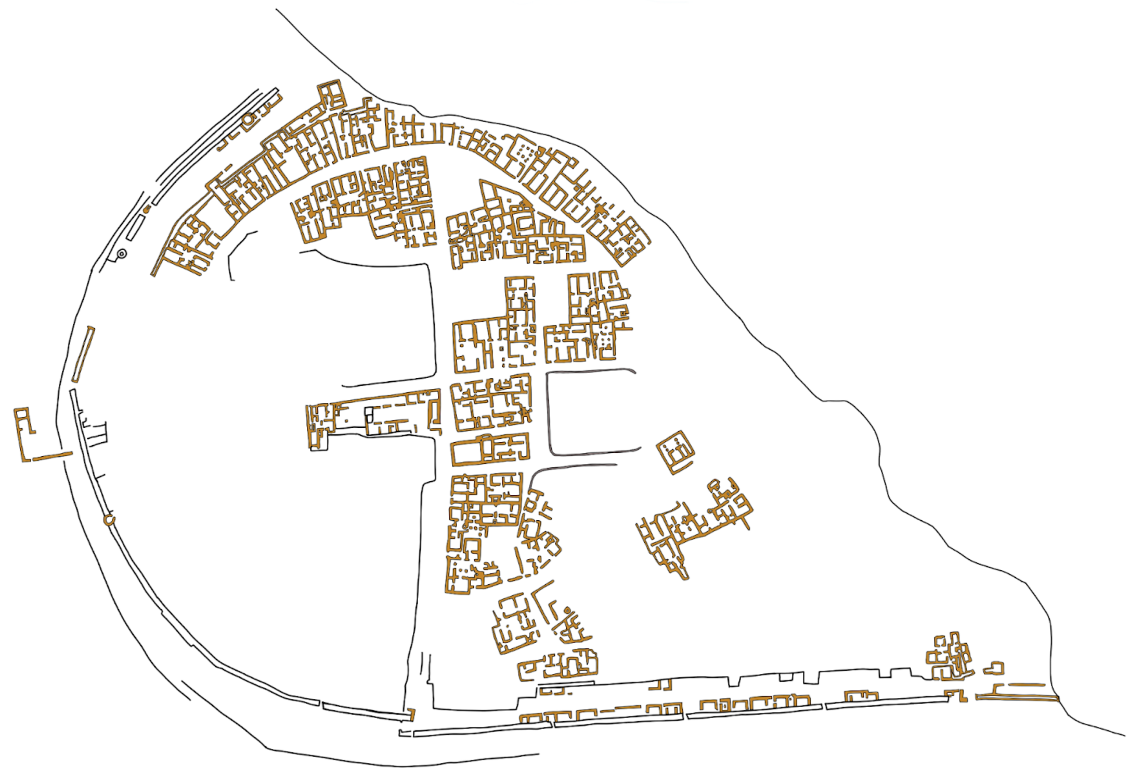

**Fig. S16. Kerkouane site map**

The excavated portions of Kerkouane, based on (Fantar 1984).

### Supplementary Tables

#### Table S1. Runs of homozygosity

| Sample | Site | Number of ROH segments (>5Mb) | Total length of ROH | Length of longest ROH segment |
|---|---|---|---|---|
| **R11778.SG** | Kerkouane | 4 | 23 | 8.5 |
| **R10339.SG** | Tarquinia | 5 | 25 | 6 |
| **R11753.SG** | Kerkouane | 19 | 424.9 | 57.2 |
| **R11791.SG** | Kerkouane | 24 | 391.3 | 53.6 |

Summary of homozygosity for individuals with more than one homozygosity segment (>5Mb).

### Supplementary Datasets

#### Dataset S1. Newly reported ancient individuals

Information about the newly reported individual genomes in the study, including sample ID, burial number, dates, latitude, longitude, site, country, genome-wide coverage, SNP coverage, and contamination estimates.

#### Dataset S2. Ancient individual genomes used in analyses

The IDs and populations categories for previously published ancient genomes used in the study. Each tab in the dataset shows the individuals and populations used for a different set of analyses. Tab 1 shows the individuals used in PCA. Tab 2 shows the individuals used in Admixtools (qpWave and qpAdm) analyses. Tab 3 shows the source populations used in supervised ADMIXTURE analysis.

#### Dataset S3. AMS dating and isotope analysis results

AMS dating and δ13C (carbon) and δ15N (nitrogen) isotopic results for the newly reported individuals in this study. The results were calibrated using the intCal20 calibration curve using the OxCal interface (https://c14.arch.ox.ac.uk/oxcal/OxCal.html).

#### Dataset 4. Admixture and cladistic analysis using qpAdm and qpWave

Tables reporting the admixture proportions, p-values, chi-squared statistic, and standard errors from qpAdm and pair-wise qpWave modeling.

#### Data availability

All tools and data needed to reproduce and evaluate the conclusions in this paper are presented in the main text and the Supplementary Materials. Alignment files for the DNA sequences for all newly reported individual genomes will be available at the European Nucleotide Archive (ENA) database under the accession number Project PRJEB49419. (www.ebi.ac.uk/ena/browser/view/PRJEB49419).

34. Maier, R. *admixtools*. (Github).

35. Olalde, I. *et al.* The genomic history of the Iberian Peninsula over the past 8000 years.

*Science* **363**, 1230–1234 (2019).

36. Fernandes, D. M. *et al.* The spread of steppe and Iranian-related ancestry in the islands of the western Mediterranean. *Nat Ecol Evol* **4**, 334–345 (2020).
37. Alexander, D. H., Novembre, J. & Lange, K. Fast model-based estimation of ancestry in unrelated individuals. *Genome Res.* **19**, 1655–1664 (2009).
38. Fregel, R. *et al.* Ancient genomes from North Africa evidence prehistoric migrations to the Maghreb from both the Levant and Europe. *Proc. Natl. Acad. Sci. U. S. A.* **115**, 6774–6779 (2018).
39. van de Loosdrecht, M. *et al.* Pleistocene North African genomes link Near Eastern and sub-Saharan African human populations. *Science* **360**, 548–552 (2018).
40. Rodríguez-Varela, R. *et al.* Genomic Analyses of Pre-European Conquest Human Remains from the Canary Islands Reveal Close Affinity to Modern North Africans. *Current Biology* vol. 27 3396–3402.e5 Preprint at https://doi.org/10.1016/j.cub.2017.09.059 (2017).
41. Arauna, L. R. *et al.* Recent Historical Migrations Have Shaped the Gene Pool of Arabs and Berbers in North Africa. *Mol. Biol. Evol.* **34**, 318–329 (2017).
42. Schuenemann, V. J. *et al.* Ancient Egyptian mummy genomes suggest an increase of Sub-Saharan African ancestry in post-Roman periods. *Nat. Commun.* **8**, 15694 (2017).
43. Fadhlaoui-Zid, K. *et al.* Genome-wide and paternal diversity reveal a recent origin of human populations in North Africa. *PLoS One* **8**, e80293 (2013).
44. Lucas-Sánchez, M., Serradell, J. M. & Comas, D. Population history of North Africa based on modern and ancient genomes. *Hum. Mol. Genet.* (2020) doi:10.1093/hmg/ddaa261.
45. Henn, B. M. *et al.* Genomic ancestry of North Africans supports back-to-Africa migrations. *PLoS Genet.* **8**, e1002397 (2012).
46. Fregel, R. *et al.* Mitogenomes illuminate the origin and migration patterns of the indigenous people of the Canary Islands. *PLoS One* **14**, e0209125 (2019).
47. Marcus, J. H. *et al.* Genetic history from the Middle Neolithic to present on the

Mediterranean island of Sardinia. *Nat. Commun.* **11**, 939 (2020).
